# A retinal circuit that vetoes optokinetic responses to fast visual motion

**DOI:** 10.1101/2021.10.31.466688

**Authors:** Adam Mani, Xinzhu Yang, Tiffany Zhao, Megan L. Leyrer, Daniel Schreck, David M. Berson

## Abstract

Optokinetic nystagmus (OKN) complements the vestibulo-ocular reflex (VOR) to stabilize the retinal image during head rotation. OKN is driven by the ON direction-selective ganglion cells (ON DSGCs), a rare class of retinal output neuron that encodes both the direction and speed of global retinal slip. The cells and synaptic circuits that give ON DSGCs their directional tuning are well known, but those dictating their slow-speed preference (and thus OKN’s) remain enigmatic. Here, we probe this circuit through patch recordings, functional imaging, genetic manipulation, and serial electron microscopic reconstruction in mouse retina. We confirm earlier evidence that feedforward glycinergic inhibition is the main suppressor of ON DSGC responses to fast motion and reveal a surprising source for this inhibition ─ the VGluT3 amacrine cell, a retinal interneuron that releases both glycine and glutamate, exciting some neurons and inhibiting others. We find that VGluT3 cells respond robustly to fast global motion and that their output reaches most RGC types, as well as a diverse group of amacrine and bipolar cells. They *enhance* the response of ON-OFF DSGCs to fast motion, while *suppressing* it in ON DSGCs. Together, our results identify a novel role for VGluT3 cells, limiting the range of retinal slip speeds that drive image-stabilizing eye movements. More broadly, they suggest VGluT3 cells shape the response of many RGCs and amacrine cells to fast motion.

## Introduction

Animals see better when their visual systems are supported by sensorimotor networks for image stabilization. Stabilizing the image on the photoreceptor array simplifies crucial visual computations, such as extracting weak, noisy signals or distinguishing the motion of objects from retinal image motion resulting from the observer’s own eye, head, or body movements.

Image-stabilizing reflexes are extremely widespread among animals with image-forming eyes. In vertebrates, rotation of the head triggers vestibular and visual feedback signals to brainstem and cerebellum oculomotor networks which generate eye counter-rotation to cancel the global drift of the retinal image (retinal slip). These innate responses comprise the vestibulo-ocular reflex (VOR) and the optokinetic reflex (OKR) [1–3]. Both produce nystagmus, in which slow image-stabilizing eye movements are periodically interrupted by fast resets of gaze in the opposite direction.

Vestibular and visual image-stabilizing reflexes complement one another in the velocity domain. The VOR plays the primary image-stabilizing role during rapid rotatory motion of the head. Canal signals are strong, driving a powerful VOR that nulls most of the retinal image motion. The modest residual slip triggers the OKR. At slow speeds, for which VOR gain is weak, OKR is the dominant stabilization mechanism and the two reflexes sum to achieve nearly perfect stabilization over a broad range of rotational speeds. The OKR is speed-tuned. It must be responsive to slow slip velocities to compensate for the limitations of the VOR. but it must be insensitive to fast speeds or it would thwart fast eye movements, such as saccades or the fast resetting phase of nystagmus.

Among the more than three dozen types of mammalian retinal ganglion cells (RGCs) [4–6], only a very specific subset appear to participate in visually driven image stabilization. These are direction-selective ganglion cells (DSGCs), and mainly those of a specific subclass – the ON DSGCs. They send their axons almost exclusively to the nuclei of the accessory optic system (AOS) [1,2,7], which relays their retinal slip signals to the vestibulocerebellum and through it to the oculomotor system. This circuit, including apparently homologous ON DSGCs, is highly conserved across vertebrates. The far more common ON-OFF class of DSGCs makes only a small contribution to this circuit, projecting instead mainly to the lateral geniculate nucleus and superior colliculus.

ON DSGCs, like the OKR they drive, respond best to slow speeds of retinal slip, typically ∼1°/s, depending on species and stimulus [2,8,9]. They are effectively silent during fast visual motion, as during saccades or the fast phase of nystagmus. By contrast, ON-OFF DSGCs respond well to such fast motion [9] (but see [10]). Other types of RGCs exhibit diverse speed tuning profiles, with some highly responsive to very fast motion.

ON DSGCs thus encode both components of the vector of retinal slip used by the image-stabilization system – its speed and direction. The mechanisms conferring direction selectivity on these and other DSGCs, anchored by starburst amacrine cells (SACs), have been studied in great detail [11–14]. By contrast, much less is understood about the cell types, neurotransmitters, and synaptic circuits that confer slow speed tuning upon ON DSGCs or distinct speed-tuning profiles upon other RGC types.

Diverse mechanisms may underlie slow-speed tuning in ON DSGCs. Excitatory drive might be attenuated at higher speeds due to suboptimal spatiotemporal summation of excitatory inputs across the dendritic arbor [15] or intrinsic membrane properties that filter out the high-frequency excitatory currents induced by fast motion. Alternatively, fast motion might trigger presynaptic inhibition of the bipolar cells (and VGluT3 amacrine cells [16]) which provide excitatory input to ON DSGCs.

In an alternative candidate mechanism, amacrine cells activated by fast global retinal slip would provide direct feedforward inhibition onto the ON DSGC. A study in rabbit retina showed that just such a circuit confers slow speed tuning on ON DSGCs, and demonstrated that the inhibitory transmitter is glycine [9]. The closely related ON-OFF DSGCs lack this glycinergic input and respond well to fast speeds [9]. Here in mouse retina we have used serial block face electron microscopy (SEM), patch recordings, and cell-type-specific optogenetics and chemogenetics to delineate the synaptic networks responsible for vetoing ON DSGC responses at high speeds. Our observations in mice echo the rabbit findings: direct feedforward glycinergic inhibition of ON DSGCs is primarily responsible [9] for their slow speed preference.

We next exploit connectomics to identify an unexpected source of this glycinergic inhibition ─ the VGluT3 amacrine cell ─ an unusual amacrine cell that makes excitatory (glutamatergic) synapses onto some postsynaptic targets and inhibitory (glycinergic) synapses onto others [17–19]. A prior optogenetic study found an excitatory glutamatergic VGluT3 input to ON DSGCs [16]. Though we replicate that result in a few ON DSGCs, the dominant synaptic influence is glycinergic inhibition. We demonstrate that this provides the main basis for vetoing the ON DSGCs response to fast global retinal slip.

Previous studies have shown that VGluT3 cells have strong receptive-field surrounds [16, 20], which would seem incompatible with robust responses to fast global motion. However, our calcium imaging data demonstrate robust activation of VGluT3 dendrites by global motion, weakened surrounds when the stimulus is moving, and a marked preference for fast motion.

We confirm that the other major class of DSGCs, the ON-OFF DSGCs ─ receive only excitatory glutamatergic input from VGluT3 cells. This *augments* their responses to fast motion, in contrast to the suppressive influence in ON DSGCs. The connectomic data document widespread VGluT3 output to most types of RGCs. This suggests that these amacrine cells may play a major role in sculpting the retinal output during fast motion, by enhancing the responses of some RGCs and inhibiting others. In ON DSGCs, it vetoes the response of the image-stabilization system to fast velocities such as those generated by gaze shifts, neatly separating these two visuo-oculomotor realms.

## Results

### Inhibition at fast speeds underlies slow speed tuning in mouse ON DSGCs

To explore the speed-tuning mechanism, we made patch recordings of synaptic currents and spiking responses in ON DSGC during presentation of full-field gratings drifting in the preferred direction at various speeds. We located these rare RGCs using 2-photon imaging of fluorescent reporters (Hoxd10-GFP or Pcdh9-Cre) or by surveying extracellular spike responses of RGCs of intermediate size for characteristic responses to flashed stimuli (see Supplemental Information and Methods). In all cases, we confirmed the identity of ON DSGCs from their selectivity for the direction of grating drift and through dye-filling and imaging of their characteristic dendritic morphology (see Supplemental Information and Supp. Fig. S1).

As expected, ON DSGCs were remarkable among RGCs in preferring slower speeds of grating drift (Fig. 1A-C, blue curves). Optimal spiking responses were obtained at retinal speeds of 150 ± 11 µm/s (angular velocity of 5 ± 2°/s; mean ± SEM; n = 63), close to previously reported values for mouse ON-DSGCs and OKN gain [2]. This is probably a slight overestimate of preferred speed; about one in four cells responded best to the slowest speed tested (76 µm/s) and thus may have preferred even slower speeds. Spike responses were strongly attenuated at speeds above the optimum, dropping to half their maximum spike rate at roughly twice the optimum speed (360 ± 23 µm/s, 12 ± 0.8°/s). Many other RGC types responded to much higher speeds of grating drift (Fig. 1C). For example, ON-OFF DSGCs exhibited optimum speeds about four times faster than those of ON DSGCs (600 ± 70 µm/s; half max. speeds 2000 ± 160 µm/s, 67 ± 5 °/s, (n = 17), Fig. 1C). Some RGC types responded to even higher speeds. For example, ON alpha cells maintained better than half-maximal responses even above 3000 µm/s (100 °/s) . Other RGC types, such as ON-delayed RGCs [21], shared the slow speed tuning of ON DSGCs.

**Figure 1.**
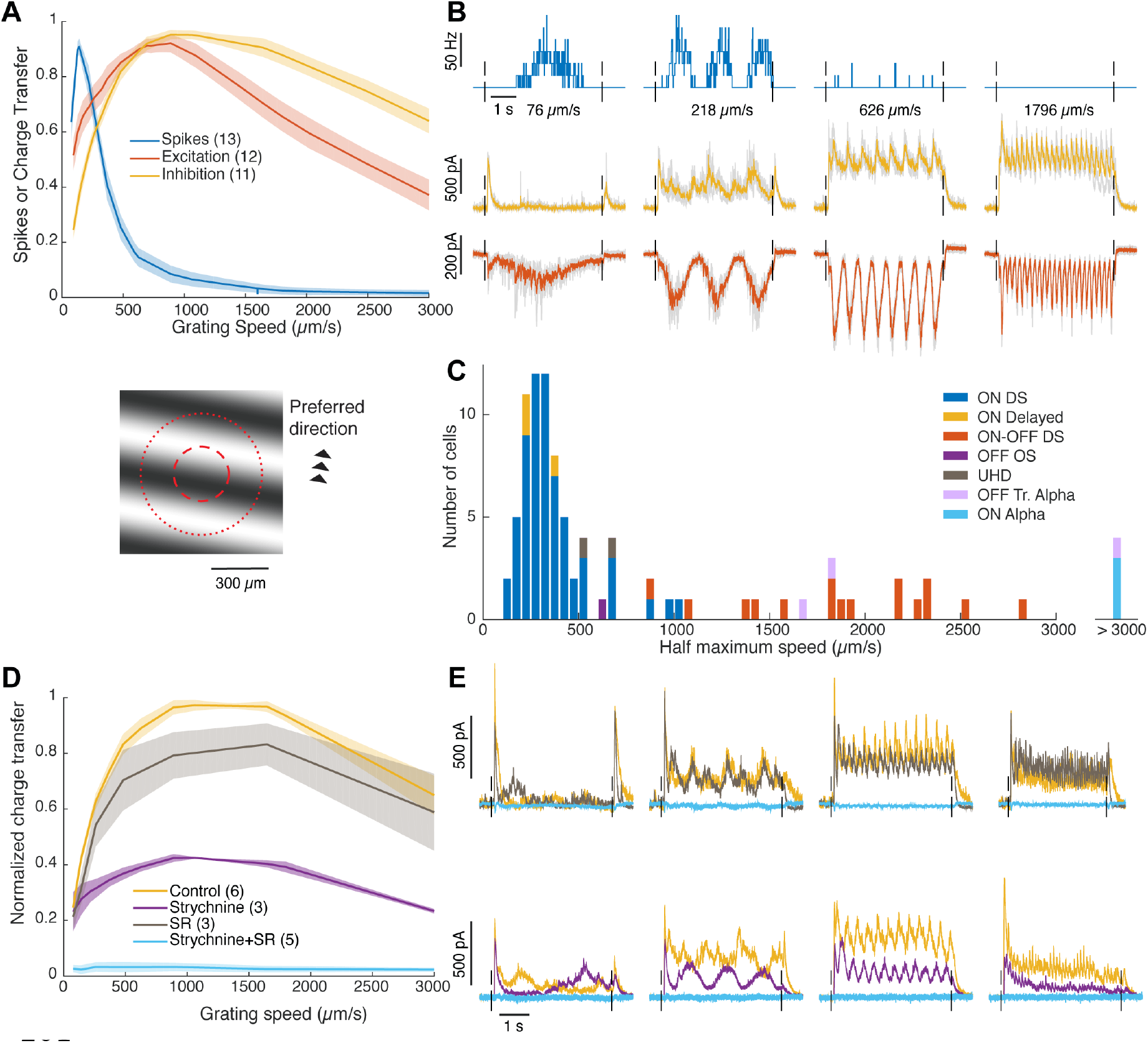
Inhibitory mechanism underlying slow speed tuning in ON DSGCs., (A) Top: Speed tuning in the response of ON DSGCs to moving gratings, as revealed by spiking (blue) and synaptic currents (excitation: red; inhibition: orange). Curves are population averages; all are normalized to their maxima. Shading denotes standard error of the mean (S.E.M.). Currents, expressed as charge transfer, were measured at a holding voltage of -65 mV (excitation) and +20 mV (inhibition). Bottom: the grating stimulus shown to scale with a schematic ON DSGC receptive field (center and surround; red circles). (B) Time course of mean spiking and current responses (±S.E.M.) to different grating speeds; color scheme as in A. (C) Distribution of speed preferences among RGCs of different functional types as indexed by the highest speed capable of evoking a response half that evoked by the optimal speed. (D) Pharmacological analysis of the inhibitory conductance. Selective glycinergic blockade (purple; strychnine) strongly reduces the inhibition seen under control conditions (orange), while blocking ionotropic GABA receptors (black; SR95531) had a weaker effect. Combined application (blue) blocked all inhibition. (E) Effects of GABA blockade (black; top row) or glycinergic blockade (purple; bottom row) on the time course of inhibitory currents. In both cases, the other antagonist was subsequently added to the first, blocking all inhibition (blue).

We hypothesized that the preference of mouse ON-DSGCs for slow speeds would result from direct glycinergic inhibition, as demonstrated in rabbit retina [9]. To test this idea, we first used whole-cell voltage clamp to record excitatory and inhibitory currents evoked in ON DSGCs by various speeds of grating drift in the preferred direction (Fig. 1A,B, red, orange curves). Both excitatory and inhibitory input were broadly tuned for speed, but with different profiles. Excitation was present at the slowest speeds we tested, but rose with speed, peaking at speeds five times faster than those evoking the most robust spiking (710 ± 90 µm/s (n = 11), or 24 °/s).

Excitation declined to half its maximum at 2290 ± 170 µm/s. Thus, although excitatory drive is affected by speed, this is insufficient to explain the slow speed tuning in ON DSGCs.

Though inhibition was strongest at speeds close to those evoking the strongest excitation (1220 ± 170 µm/s, declining half max. > 2900 µm/s, n = 12), it was relatively weaker than excitation at slow speeds and relatively stronger at fast ones. The crossover of the two curves occurred at ∼700 µm/s, above which nearly all spiking was suppressed (Fig. 1A). Taken together, the above results imply that slow speed tuning of ON DSGCs reflects a shift in excitatory-inhibitory balance in favor of inhibition at higher speeds, as in the rabbit [9].

For most speeds, the excitatory current traces exhibited a single prominent peak for each cycle of the grating (Fig. 1B), presumably reflecting glutamatergic input from ON bipolar cells, their dominant source of excitation. In contrast, inhibitory current traces often exhibited minor peaks roughly anti-phase to the excitation, when the dark bars of the grating overlie the receptive field. The OFF pathway may therefore make a larger contribution to inhibition than to excitation. At higher speeds, the inhibitory current exhibited not only the stimulus-locked modulatory component but also an underlying continuous inhibitory conductance. This was much less apparent for excitation. This confirms earlier observations in rabbit ON DSGCs.

### Fast retinal slip triggers feedforward glycinergic inhibition of ON DSGCs

We repeated these experiments adding receptor antagonists to the bath to test the hypothesis that the feed-forward inhibition evoked at fast speeds was glycine-mediated, as in rabbit (Fig. 1D,E). The glycine receptor antagonist strychnine sharply attenuated the inhibition, dramatically reducing the steady plateau of inhibition evoked by fast grating motion. At speeds evoking maximum inhibitory current, total inhibition was reduced by more than half its control value (57±4% decrease in charge transfer; n=3. See note in Supplemental Information). In contrast, blockade of GABA_A_ receptors alone (SR95531) had only a modest effect (Fig. 1D,E; 17±7% decrease in charge transfer; n=3). Simultaneous application of both antagonists abolished all inhibitory current in the ON DSGCs. Thus, effectively all inhibition driven by fast motion is mediated by GABA_A_ and glycine receptors under our experimental conditions.

Taken together, these results demonstrate that fast motion triggers direct glycinergic inhibition of ON DSGCs, as first shown in rabbit [9]

### Electron microscopic reconstructions identify VGluT3 amacrine cells as the primary source of glycinergic synapses onto ON DSGCs

Most glycinergic retinal inhibition derives from small-field amacrine cells with highly branched dendritic arbors [22, 23]. We searched for synaptic contacts from such cells onto ON DSGCs in an existing serial blockface electron microscopic (SEM) dataset of the adult mouse inner plexiform layer spanning >200 μm (volume ‘k0725’ [24]). We focused our analysis mainly on a single previously reconstructed presumed ON DSGC (Fig. 2) [15]. Numerous anatomical features identify this as an ON DSGC. Its dendrites narrowly costratify with the processes of starburst amacrine cells (SACs), with nearly the entire arbor within the ON SAC plexus (Fig. 2B-E). There, they cofasciculate with SAC processes (Fig. 2A) and receive numerous wrap-around synaptic contacts [12] from SAC varicosities (Fig. 2F, G). Analysis of the orientation of the presynaptic SAC processes confirmed the asymmetric connectivity that confers direction selectivity upon DSGCs [12–15]. The minor OFF branches, a common feature of ON DSGC arbors in mice [2], receive some OFF SAC contacts. Presumptive ON-OFF DSGCs in the volume shared many of these attributes but had smaller dendritic arbors, higher branching density, and a larger fraction of their arbor in the OFF SAC plexus.

**Figure 2.**
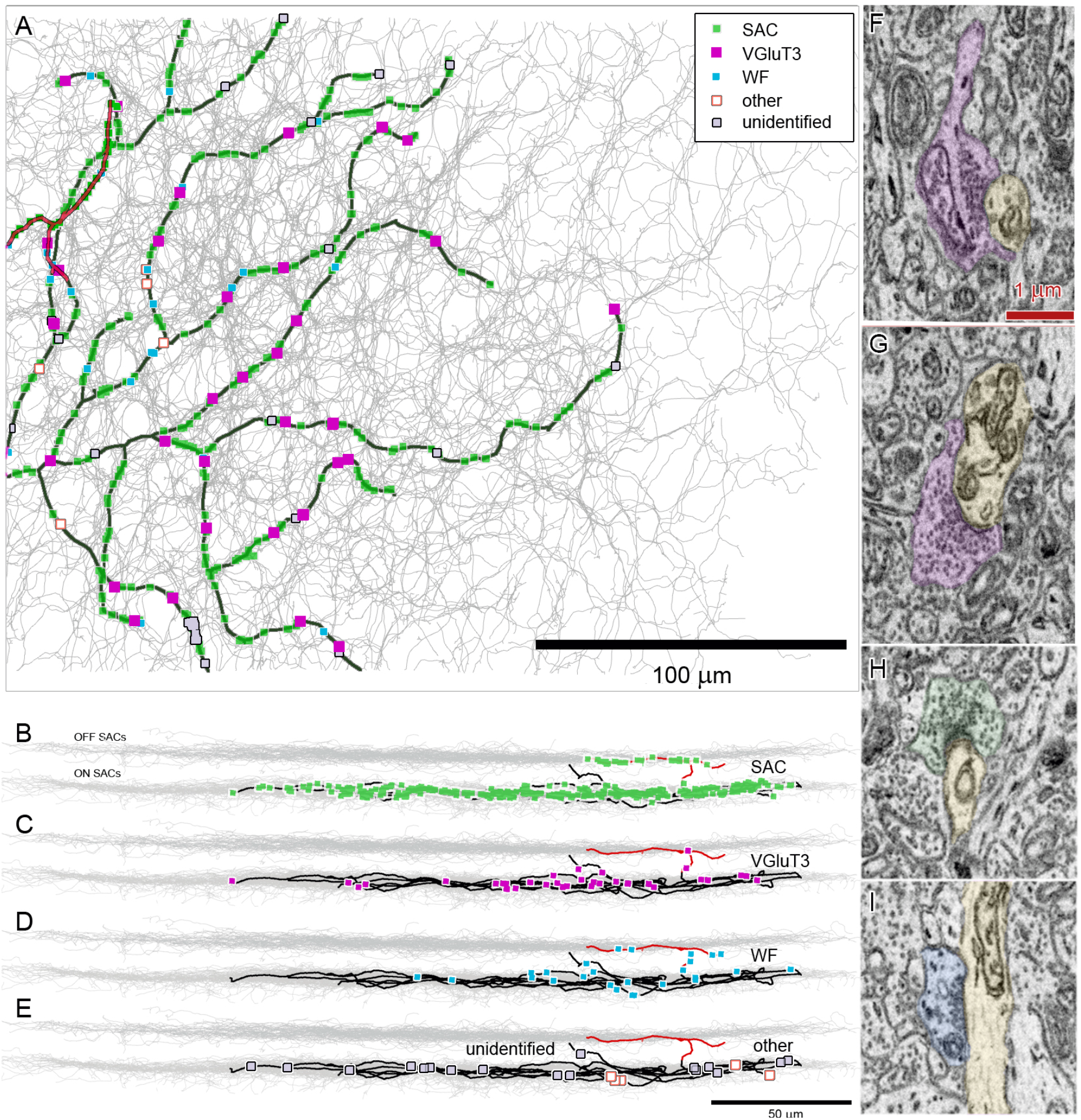
Amacrine cell inputs onto an ON DSGC. (A) SEM reconstruction of the distribution and sources of all detected amacrine-cell synaptic contacts onto an ON DSGC. Dendrites of the ganglion cell are shown in black except for the sparse branches in the OFF sublayer (red). Synaptic contacts are plotted as dots, color coded by amacrine-cell source (see key), as determined by reconstruction of presynaptic cells. The plexus of ON SAC dendrites is shown in gray. Boundaries of the SEM volume appear as a fine rectangle. (B-E) side (vertical) views of depth of these inputs within the IPL. Gray bands mark the ON and OFF SAC plexuses. (G-I) Example SEM images of amacrine-cell synapses onto the ON DSGC. The ON-DSGC is shown in yellow. The presynaptic cells are color coded as in A-E: VGluT3 cells (F,G), an ON SAC (H) and a WF amacrine cell (I).

To identify possible sources of glycinergic amacrine cell inhibition to this ON DSGC, we first marked all the non-ribbon synaptic contacts we could find (n=431; Fig. 2F-I). For each contact, we reconstructed enough of the presynaptic process to sort it into one of three groups (Fig. 2A-E): SACs; wide-field amacrine cells; or candidate glycinergic neurons. SACs were recognizable from their narrow stratification within the ON or OFF SAC plexus, thin straight connecting dendrites, and large wrap-around varicosities targeting mainly other SACs and DSGCs. Wide-field cells had sparser branching and fewer varicosities and were not strictly confined to the SAC plexuses. Candidate glycinergic neurons were much more highly branched, with arbors that extended well outside the SAC plexuses.

SAC inputs accounted for fully 83% of the non-ribbon contacts (n=340; 317 ON and 23 OFF). At least by this measure, GABAergic SAC input dominates the inhibition of ON DSGCs. Candidate glycinergic inputs (n=41) were the next most common, comprising 10% of non-ribbon inputs, with wide-field inputs nearly as common at 7% of all input (n=27). A small minority of inputs (n=23; 5%) could not be identified because too small a fraction of the arbor could be reconstructed, and were excluded from the sample.

We extensively reconstructed all of the presumptive glycinergic neurons identified by our initial screen. To our surprise, nearly all of them appeared to be VGluT3 amacrine cells, as documented below (n=38 synapses from 16 cells). Two were synapses from Type H18 amacrine cells [25, 26] and one was from an unidentified medium-field type. We detected no synaptic inputs from any of the many other types of small-field amacrine cell types, even after examining a large sample of such cells reconstructed in another study [26]. We conclude that VGluT3 amacrine cells are by far the best candidate glycinergic cell type shaping the slow-speed tuning of ON DSGCs.

Multiple structural observations confirm these as VGluT3 amacrine cells (Fig. 3). First, their stratification is appropriate (Fig. 3E, F, J). They arborize most heavily in the middle of the inner plexiform layer (IPL), between the ON and OFF SAC plexuses, but extend sparse processes into and even a bit beyond those plexuses . Their cell bodies lie in the inner nuclear layer, are of appropriate size, and are distributed with spacing that suggests a regular mosaic (Fig. 3A). Their dendritic-field diameters (106 ± 7 μm, n = 10 cells) are in line with earlier data [18, 20] and overlap extensively with a coverage factor of at least 5 (Fig. 3A, inset).

**Figure 3.**
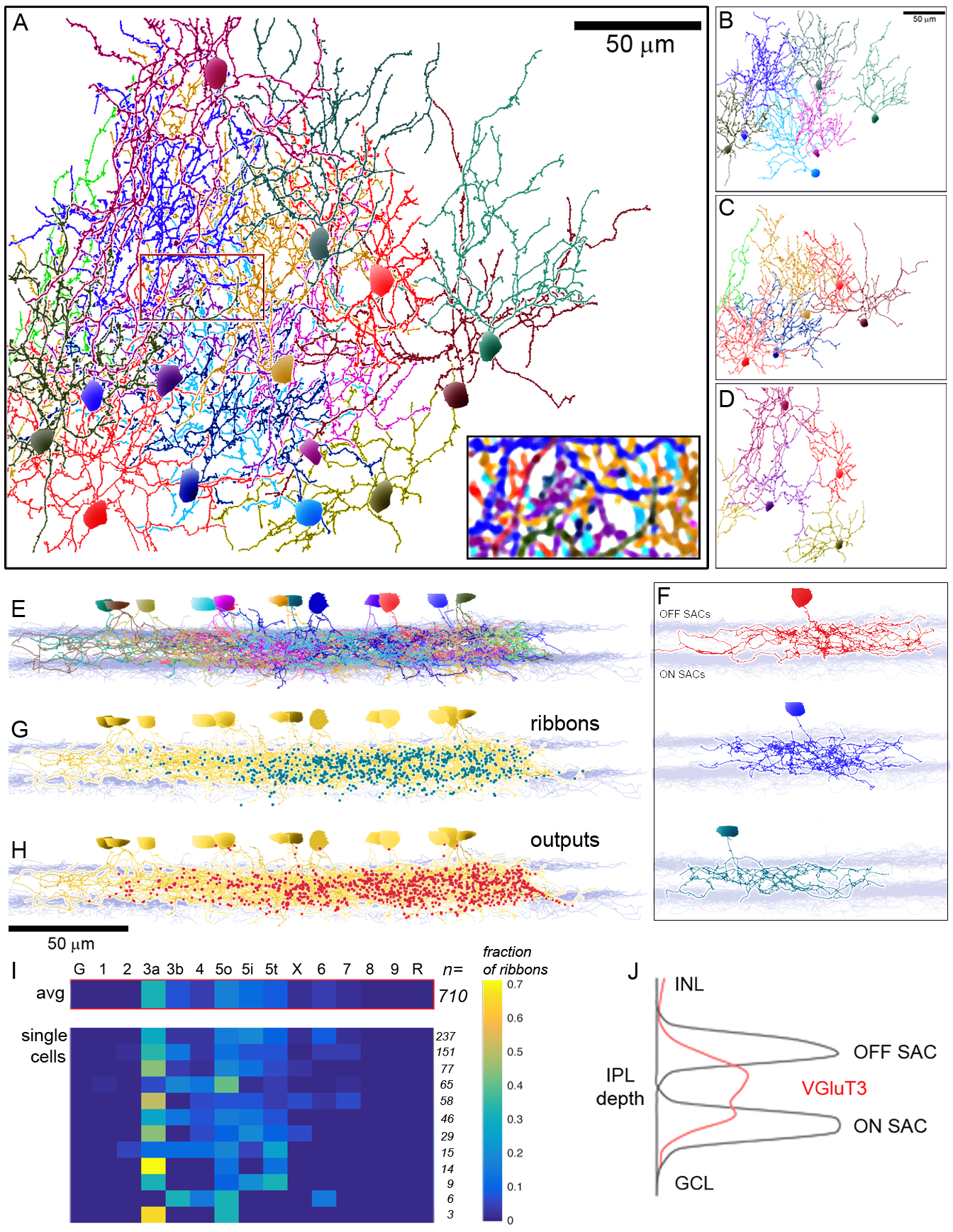
Morphology, input and output of VGluT3 amacrine cells. (A-D). Dendritic arbors and mosaic of VGluT3 amacrine cells reconstructed in the k0725 SEM volume as seen in top view (*en fac*e). Enlarged view (inset) shows that local regions contain dendrites of at least 5 VGluT3 cells. B-D: Same dendritic profiles as in A but divided arbitrarily into three groups to provide a clearer view of single arbors. (E,F) Side (vertical) views of the stratification of reconstructed VGluT3 cells within the IPL, shown together (E) and, for 3 cells, individually (F). (G, H) Side view of the location in depth of ribbon inputs (F) and synaptic outputs (G) of VGluT3 cells. Gray bands in E-H mark the ON and OFF SAC plexuses; VGluT3 cells appear gold in G and H. (I) Heatmap plot of the fraction of all ribbon synaptic inputs derived from each bipolar type, identified at the top (G – GluMI, X – XBC, R – rod bipolar cell). Bar at top shows pooled data for all identified ribbon contacts (n=710). Lower panel illustrates the consistency of this pattern across individual cells, shown one per row in descending order of the number of identified ribbons. (J) Stratification profile of reconstructed VGluT3 processes (red; n=17) in relation to those of ON and OFF SACs (black). Data are normalized to the ON SAC plexus.

Input and output patterns matched previous reports on VGluT3 cells, and were very consistent across the population of reconstructed cells (Fig 3I, Supplemental Table S5), confirming other evidence that they belong to a single type. In keeping with functional evidence for input from both the ON and OFF pathways to VGluT3 cells, our reconstructed cells receive ribbon synapses from diverse ON and OFF bipolar cells.

The complement of bipolar inputs largely reflects their dendritic stratification (Fig. 3G, I) with most input from OFF types 3a, 3b, and 4 and ON types 5o, 5i, 5t, and little from bipolar types stratifying at the IPL margins (OFF types 1,2 and GluMI [27]), and ON types 6, 7, 8 and 9. Their outputs include all the ganglion-cell targets so far identified in physiological studies (Supplemental Table S5). These included W3 cells [16, 28], OFF transient alpha cells [29], Suppressed-by-Contrast cells of several types [29, 30] including the ON Delayed RGC [21] (see Discussion), both ON and ON-OFF DSGCs [16, 28], and M1 ipRGCs [31] (For RGC type nomenclature see [4] and accompanying online database). However, our SEM analysis suggests that their outputs are much more diverse than previously appreciated, with contacts to the great majority of known RGC types [4], including OFF sustained alpha, Jam-B [32], ON alpha, F-mini-ON [33], F-mini-OFF [33], OFF transient medium RF, and diverse varieties of smaller field RGCs stratifying between the ChAT bands [34]. Two ipRGC types notably not targeted by VGluT3 are the M2 [35] and M5 (PixON) [36, 37] types, which stratify largely outside the VGluT3 plexus. VGluT3 cells also synapsed upon diverse types of amacrine cells, though only rarely onto other VGluT3 cells or SACs.

We have not reconstructed most amacrine-cell targets, though WF cells stratifying between the SAC plexuses were among the most common recipients [38]. VGluT3 outputs, like their ribbon inputs, were distributed most heavily within and between the SAC plexuses (Fig. 3H), though they were also sparsely present in the distal IPL. Some of these are somato-dendritic synapses onto M1 ipRGCs. One unexpected feature of the VGluT3 cells in this volume was a consistent upward (roughly dorsal) displacement of the dendritic arbor relative to the soma (Fig. 3B-D).

### VGluT3 cells make functional glycinergic synapses onto ON DSGCs

Of the three types of amacrine cells shown by the SEM analysis to provide input to ON DSGCs, only one – the VGluT3 cell – is known to be glycinergic. Thus, VGluT3 cells are apparently the source of most, if not all, the glycinergic suppression of ON DSGCs during rapid motion. The other two presynaptic types – SAC and widefield amacrine cells – use GABA as their sole inhibitory transmitter [23,39–41]. This is a surprising conclusion because VGluT3 ACs have been reported to provide glutamatergic excitation to ON DSGCs, not glycinergic inhibition [16]. To test for the inferred glycinergic influence, we recorded postsynaptic currents evoked in ON DSGCs by optogenetic depolarization of VGluT3 cells. For these studies, we crossed a novel strain of VGluT3-Cre mice with a strain that expresses channelrhodopsin2 (ChR2) in a Cre-dependent manner (Ai32) (See Methods and Supplemental Information, Supp. Fig. S2). Conventional photoreceptor-mediated light responses and cholinergic transmission were suppressed with a cocktail of synaptic blockers (‘photoreceptor block’; L-AP4, ACET, hexamethonium [16]). Optogenetic activation of VGluT3 cells evoked robust inhibitory currents in 81% of the ON DSGCs tested (13/16 cells) (Fig. 4A-C). Peak inhibitory postsynaptic currents averaged 33 ± 4 pA for single light pulses. We also delivered optogenetic pulse trains to mimic the temporal modulation of VGluT3 cells during grating motion (Fig. 4D). Evoked inhibitory currents persisted in postsynaptic ON DSGCs throughout a 5 s train over the full range of frequencies tested (1-8 Hz, corresponding to grating speeds of roughly 400 - 3000 μm/s for experiments shown in Fig. 1). At higher stimulus frequencies, IPSCs summed temporally to produce a continuous inhibitory conductance (Fig. 4D bottom traces), much as they did during rapid grating motion (Fig. 1B, yellow traces). Thus, inhibition of ON DGSCs by VGluT3 cells exhibits kinetics compatible with their proposed role in vetoing responses to fast motion.

**Figure 4.**
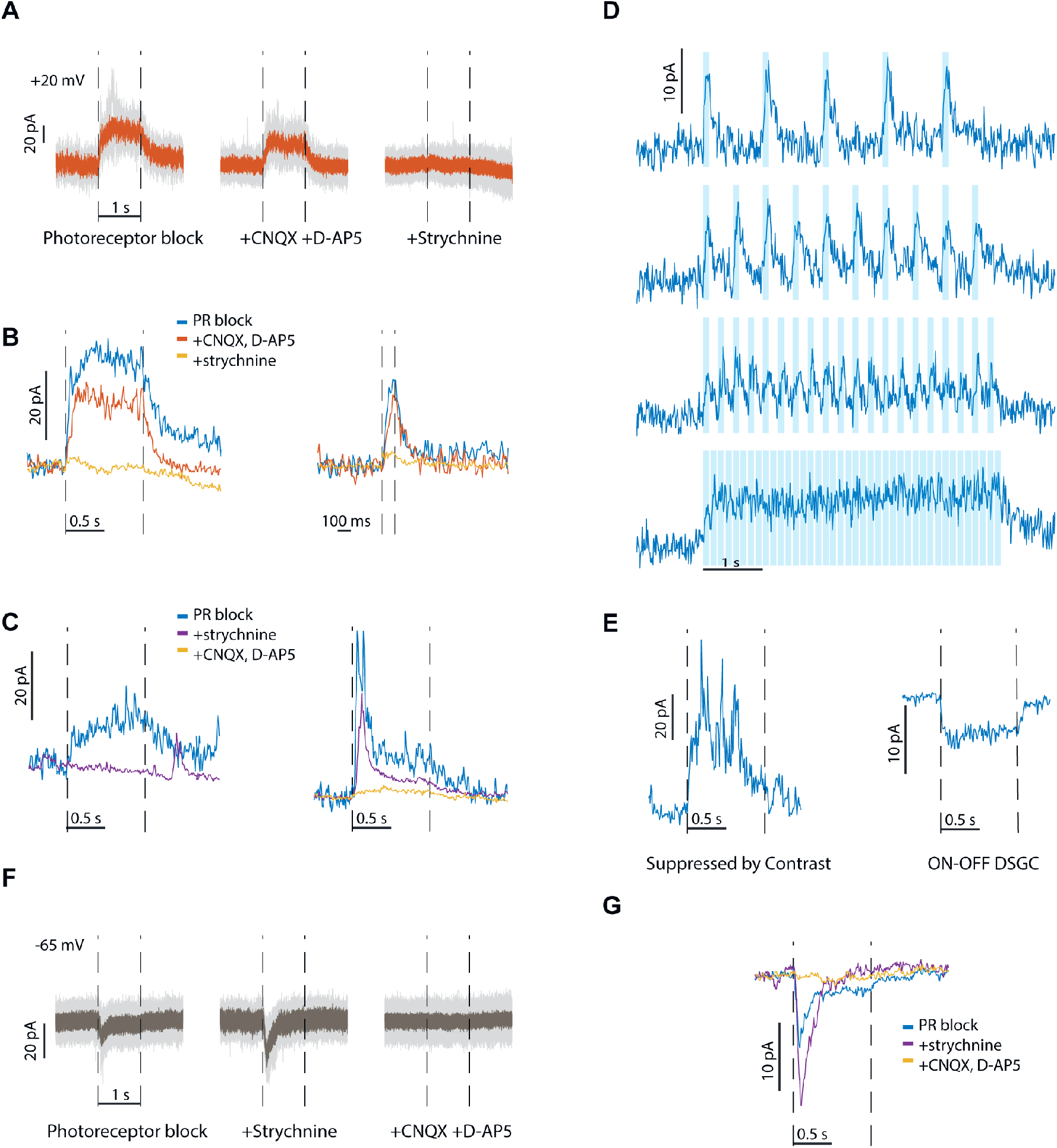
Postsynaptic currents evoked in ON DSGCs by optogenetic activation of VGluT3 cells. (A) Raw inhibitory current trace (V_hold_: +20 mV) in a single ON DSGC in response to optogenetic stimulation during the period marked by interrupted vertical lines. “Photoreceptor block” consists of L-AP4, ACET and Hexamethonium. (B) Pharmacological effects on the evoked inhibitory current in two ON DSGCs (left and right panels; left panel from same cell as in A). The sequence of drug application is listed above; current traces are smoothed. (C) As for B, but with glycinergic blockade preceding glutamatergic blockade. (D) Inhibitory currents in an ON DSGCs in response to optogenetic pulse trains of frequencies 1, 2, 4, 8 Hz. Pulse width: 100 ms. (E) Optogenetic activation of VGluT3 cells evokes inhibitory currents in a Suppressed by Contrast RGC (V_hold_: +20 mV), and excitatory currents in an ON-OFF DSGC (V_hold_: -65 mV; both under photoreceptor block). (F) Excitatory currents (raw) evoked in an ON DSGC by optogenetic activation of VGluT3 cells. V_hold_: -65 mV. (G) Pharmacological effects on this current in the same cell.

The inhibition evoked by activation of VGluT3 cells was glycinergic, as expected. Blocking ionotropic glutamate receptors (CNQX, D-AP5) had little effect on the optogenetically induced inhibitory conductance (Fig 4A, B; mean reduction in maximal current of 28 ± 7%, 5 cells), whereas further blockade of glycine receptors (strychnine) virtually eliminated the current (decrease of 81 ± 5%, 4 cells). The broad-spectrum glutamate blockade used ensures that these photostimulation-induced currents are mediated optogenetically, not because rods and cones are driving glutamatergic signals through synaptic networks. It also excludes indirect effects mediated by evoked release of glutamate from VGluT3 cells. We conclude that these amacrine cells inhibit ON DSGCs through direct glycinergic synaptic input.

We reversed the order of drug application in three cells. When strychnine was the first inhibitory blocker applied, the optogenetically evoked inhibition was completely suppressed in one cell (Fig 4C, left). However, in two other cells, strychnine left at least half the inhibition intact. This residual inhibition was abolished upon further addition of the glutamate receptor antagonists (Fig 4C, right). Presumably this strychnine-resistant inhibitory current occurs when VGluT3 activation glutamatergically excites one or more GABAergic amacrine cell types that synapse onto ON DSGCs (see Discussion).

Control experiments confirmed that off-target ChR2 expression in Müller cells was not responsible for the evoked currents in ON DSGCs; that other known targets of VGluT3 input responded as expected to the optogenetic stimulus (Fig. 4E); and that the pharmacological effects shown in Fig. 4B,C were not simply the result of rundown of optogenetic responses over trials (Supplemental Information, Supp. Fig. S3).

To evaluate a previous report that VGluT3 cells provide glutamatergic excitation to ON DSGCs [16], rather than glycinergic inhibition as found here, we repeated these experiments with ON DSGCs clamped at the chloride reversal potential.

Optogenetically evoked EPSCs were detected in only two of the 16 cells tested (12%), with peak currents of 10 ± 1 pA (Fig. 4F, G). The excitation was mediated by glutamate: it persisted following application of strychnine, but was eliminated by application of the ionotropic glutamate receptor blockade (Fig. 4G). Both of those cells also exhibited evoked IPSCs when clamped at 0 mV.

### Chemogenetic suppression of VGluT3 cells reduces fast-motion inhibition of ON DSGCs

We tested the effects of chemogenetically suppressing VGluT3 cells on the inhibition of ON DSGCs by fast motion. We used a DREADD system (Designer Receptor Exclusively Activated By Designer Drugs) in VGluT3-Cre mice, expressing the hM4Di receptor in VGluT3 cells either by intraocular injection of a Cre-dependent viral vector or by crossing VGluT3-Cre mice with Cre-dependent DREADD mice (Methods). In either case, bath-application of the DREADD ligand (CNO) markedly reduced the inhibitory currents induced by drifting gratings at all speeds (Fig. 5A, B). At speeds evoking maximal inhibition, CNO reduced inhibitory charge transfer by 43 ± 7% (n = 3) for viral experiments and 38 ± 6% (n = 3) for those based on the genetic cross. The same ligand had no effect on inhibition in an ON DSGC in a DREADD-free mouse (Fig. 5C).

**Figure 5.**
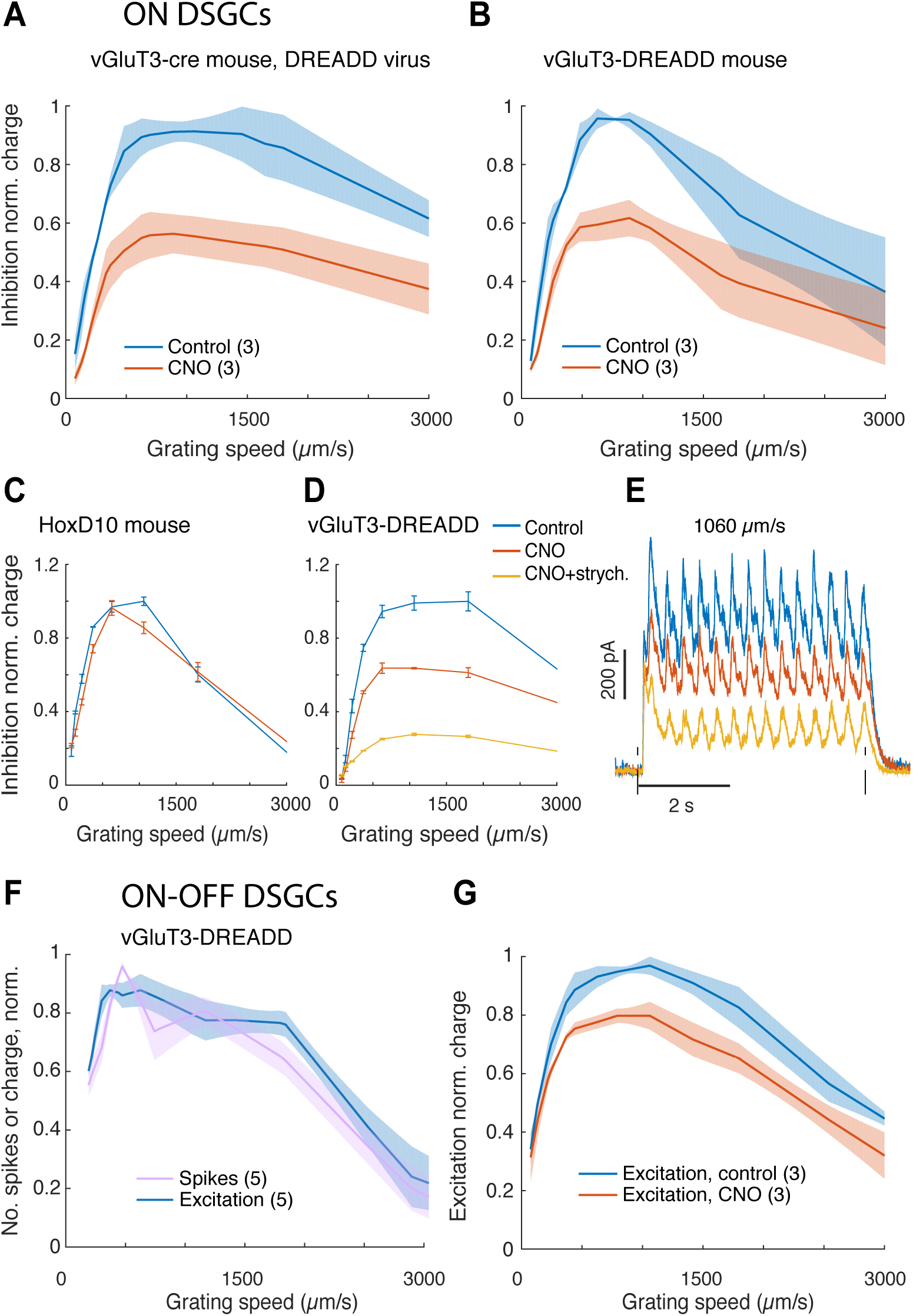
Chemogenetic suppression of VGluT3 cells alters speed tuning of ON and ON-OFF DSGCs. Effect of DREADD-mediated suppression of VGluT3 cells on speed tuning in ON DSGCs (A-E) and in ON-OFF DSGCs (F,G). (A, B) Application of the DREADD ligand CNO (red trace) partially suppressed inhibitory currents evoked by moving gratings in ON DSGCs under control conditions (blue), regardless of whether the method for Cre-dependent DREADD expression was intraocular injection of a Cre-dependent virus (A) or through a genetic cross (B). V_hold_ : +20 mV. (C) Control experiment showing that CNO has no effect when applied in a DREADD-free retina from a HoxD10-GFP mouse. (D) Most inhibition that remains after CNO application is blocked by further addition of the glycine-receptor antagonist strychnine. (E) Individual current traces of the ON DSGC in D under the same pharmacological conditions and using the same color scheme as in D. Grating speed: 1060 µm/s. (F) Spiking and synaptic excitation in ON-OFF DSGCs as a function of grating speed. V_hold_ : -65 mV (G) Speed tuning profiles of excitatory charge in ON-OFF DSGCs, before and after application of CNO. For experiments in F and G, DREADD was expressed through a genetic cross.

The chemogenetic approach (Fig. 5A, B) was less effective than strychnine (Fig. 1D) in reducing the motion-induced inhibition in ON DSGCs. Indeed, addition of strychnine during the chemogenetic manipulation further reduced the inhibition (Fig. 5D, E). This could reflect incomplete chemogenetic suppression of VGluT3 output or alternative sources of glycinergic inhibition beyond VGluT3 cells. Application of CNO scaled down the currents by about the same factor across speeds (Fig 5A, B, E) while preserving the kinetics (Fig. 5E), which is consistent with incomplete suppression of VGluT3 (see Supplemental Information and Discussion).

Taken together, the above results suggest that the native inhibition in ON DSGCs at high speeds is provided at least to a large extent by glycine release from VGluT3 cells, although our effort to suppress VGluT3 signaling was only partially effective.

### VGluT3 cells augment fast-motion responses in ON-OFF DSGCs

We found that the other direction-selective class, the ON-OFF DSGCs, were excited by optogenetic activation of VGluT3 cells excites (Fig.4E), in agreement with earlier work [16, 28]. Thus, VGluT3 cells appear to exert essentially opposite signs of effect on the two DSGC classes. ON-OFF DSGCs responded well to fast gratings that induced strong glycinergic suppression in ON DSGCs (Fig. 5F, 1C). This suggested that glutamate release from VGluT3 cells may augment ON-OFF DSGC responses in this range of speeds. Indeed, chemogenetic suppression of VGluT3 cells reduced the excitation evoked in ON-OFF DSGCs by fast gratings moving in the preferred direction (17 ± 3% reduction in excitatory charge transfer from its control value; Fig. 5G). The reduction was more pronounced at faster speeds: the speed response curves (Fig. 5G) were shifted towards lower speeds at the falling branch (Δ = 400 ± 160 µm/s at half max. of the control curves). Though incomplete blockade of VGluT3 cells may underrepresent these influences, most of the residual excitatory current presumably represents bipolar ribbon synaptic input. Thus, during rapid visual motion, VGluT3 cells affect the two DSGC classes in opposing ways, enhancing responses in ON-OFF DSGCs while suppressing ON DSGCs.

### VGluT3 dendrites are strongly activated by fast global motion

The foregoing results imply that fast motion triggers VGluT3 output. To provide more direct evidence on this point, we recorded their calcium responses using GCaMP6 expressed selectively in these cells using a genetic cross (VGluT3-Cre x Ai148) [42].

VGluT3 dendrites exhibited robust calcium signals in response to the motion of full-field grating stimuli, especially at the fast speeds that trigger feedforward glycinergic inhibition of ON DSGCs. Speed tuning was highly consistent for different ROIs within each field of view, and across most fields of view (FOV; Fig. 6C), and at different depths within the VGluT3 plexus. Some FOVs exhibited an unusually robust response to the fastest speeds (Fig. 6C, right panel).

**Figure 6.**
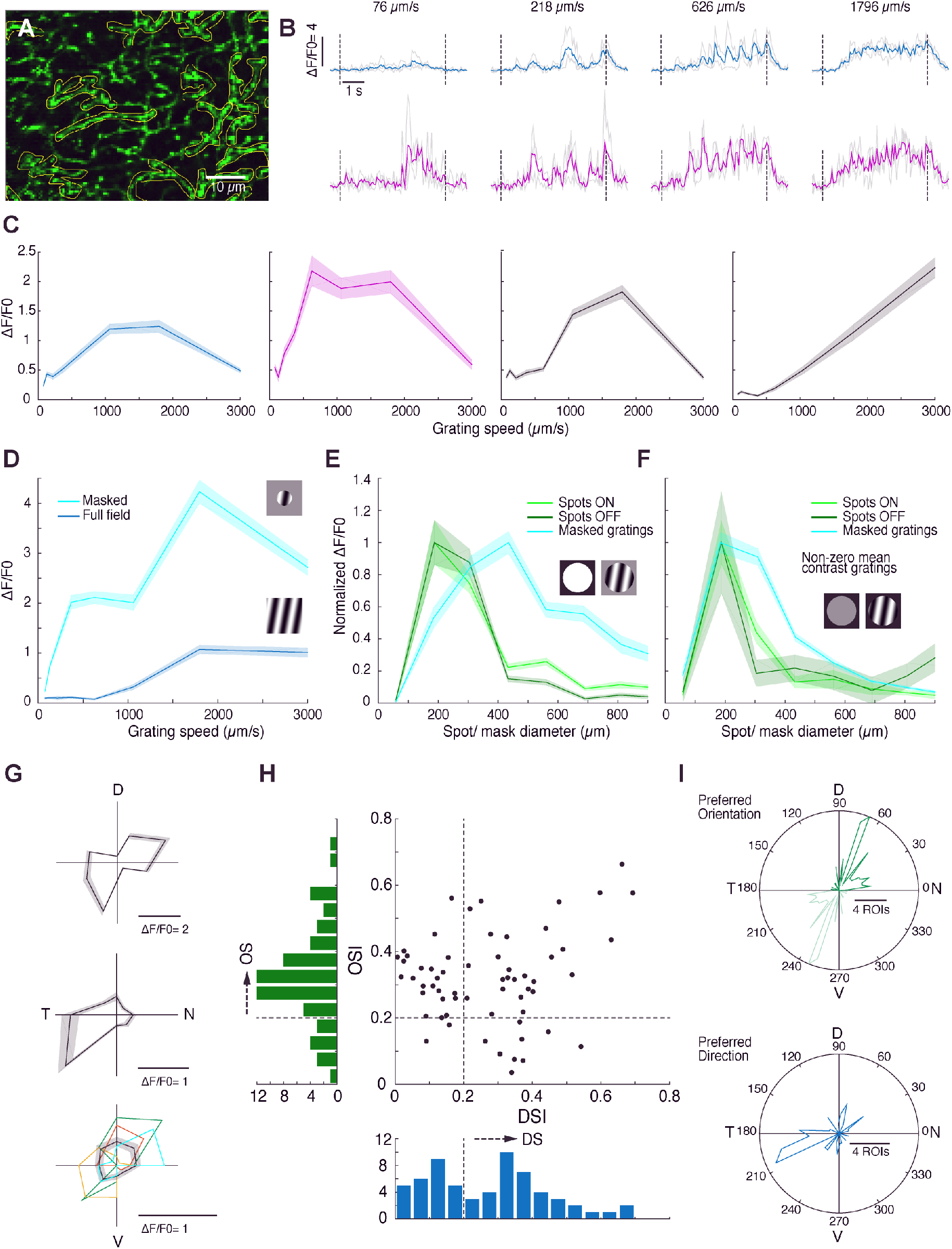
Calcium responses in VGluT3 AC dendrites. (A) Part of a field of view of a live flat-mounted retina at the level of the VGluT3 dendritic plexus, marked by GCaMP6 fluorescence. Yellow outlines show typical ROIs used to assess local Ca_2+_ signals. (B) Temporal variation in fluorescence intensity (ΔF/F_0_) in two ROIs (top and bottom rows), in response to full-field gratings drifting in one direction at different speeds, noted above the traces. Individual trials, gray; mean over trials, blue or magenta. (C) Net calcium response (ΔF/F_0_, averaged over the stimulus time) to full-field gratings as function of speed for four different FOVs; data for all ROIs in the field were pooled. The two left panels (blue and magenta) include data from the ROIs of the corresponding color in B. (D) As for C, but varying the extent of the grating to assess surround suppression of the grating response. Responses are greatly enhanced when the grating moves within a circular mask (345 µm diameter). (E) Area-response functions for flashed bright spots presented from darkness (green traces) and for drifting gratings (blue trace). Spot responses are shown separately for the ON (light green) and OFF responses (dark green). For gratings, size tuning was assessed by presentation within a circular mask of variable size (speed: 1520 µm/s; direction fixed but chosen arbitrarily, presented from gray background). (F) Same as in E., but the spots and gratings are matched in net positive contrast relative to the background. (G-I) Evidence for orientation and direction selectivity of the VGluT3 dendritic plexus. (G) Polar plots of average responses vs. the direction of motion of a full-field grating (speeds 1500 or 1900 µm/s) in 3 FOVs. Colored curves in bottom panel show data for several individual ROIs. (H) Distributions of orientation and direction selectivity indices (OSI and DSI) of 63 ROIs, 3 FOVs shown in G. An ROI was considered DS or OS if its DSI or OSI was ≥ 0.2, respectively. (I) Angular distributions of the preferred orientations (top), or preferred directions of (bottom) of single ROIS (n= 69 OS ROIs and 47 DS from 5 FOVs).

The robust response to extended stimuli was unexpected because VGluT3 receptive fields have strong suppressive surrounds [16,20,42]. Indeed, calcium responses were greatly enhanced when we restricted the same gratings to a circular patch centered on the FOV (345 µm diam.; Fig. 6D). This enhancement was apparent at all speeds, and quadrupled the response to the optimal speed. Slow gratings that failed to drive responses when presented full-field now evoked clear Ca^2+^ signals. Nonetheless, the high-speed preference remained.

To probe the spatial organization of these surround effects, we presented spots or masked gratings of various sizes centered on the FOV (Fig. 6E,F). Area-response profiles based on spot stimuli closely resembled those reported earlier based on patch recordings from VGluT3 somas [20](Fig. 6E; green curve), with strong surround suppression and maximal responses for spots ∼200 µm in diameter (ON response: 204 ±10 µm; 20 ROIs. OFF response: 217±12 µm; 19 ROIs). Grating patches (blue curve) produced a markedly different area-response profile, with much less response attenuation as grating patches extended into the surround. For the largest stimuli tested (930 µm diam.) grating responses persisted about a quarter of their maximum response (26 ± 3%, 28 ROIs), whereas spot responses were suppressed to less than a tenth of their maximum (ON: 8 ± 1%, 20 ROIs; OFF 4 ± 1%, 19 ROIs). Further, when gratings were used, the optimum size of the stimulus was roughly doubled (408 ± 10 µm; 28 ROIs). We conclude that though VGluT3 cells have strong receptive-field surrounds, the surround suppression is stimulus-dependent and permits robust responses to global retinal motion.

The difference in surround suppression between the two stimuli (Fig. 6E) could arise from the presence or absence of continuous motion. Alternatively it could be the result of a contrast increase during the stimulus time. The spot stimulus by its nature is a step in overall contrast (positive or negative) with respect to the preceding and surrounding background. The gratings however, were presented on a gray background, with the mean change in overall contrast being zero. We thus tested new sets of spots and gratings, with overall contrast in the two stimuli equal and positive (Fig. 6F; see Methods). The area-response functions for spots resembled those seen previously (Fig. 6E), though the lower contrast weakened the responses. The surround suppression for masked gratings, however, was stronger than for zero mean gratings. Despite this, receptive fields sizes assessed with gratings were once again larger than those measured with spots of equal contrast (Fig. 6F). Therefore, the weaker surround suppression in the case of gratings vs. spots, is both due to a smaller change in contrast, and due to the continuous motion.

Taken together our results suggest that VGluT3 ACs are fit to report global image slip and do not exhibit a prohibitive surround suppression.

To our surprise, the VGluT3 dendritic Ca^2+^ signals were tuned to the direction of grating motion. (Fig. 6G, H). Typically, individual ROIs exhibited a preference for grating motion in either direction along a single axis; 84% of ROIs met the standard criterion for this form of orientation selectivity (OS; 69 of 82 ROIs, 5 FOVs, 3 mice). Many ROIs preferred motion in one direction; 57% met the standard criterion for direction selectivity (DS, 47 ROIs). Only 5% (4 ROIs) showed neither orientation or direction preference (43% met both criteria). In some FOVs, ROIs were either mostly orientation selective (Fig. 6G, top) or mostly direction selective (Fig 6G, middle), while other FOVs contained a mix of OS and DS behavior (Fig 6G, bottom). Overall OS ROIs (69) were strongly clustered around a preference for dorsoventral axis of motion (∼70° or 250°); a minor second cluster appeared near the nasotemporal axis (∼20° or 200°) (Fig. 6I, left). Preferred directions of DS ROIs (47) clustered mainly near the nasotemporal axis (∼200°). Note that we defined the preferred orientation as the preferred axis of motion (not e.g. as the orientation of the bars in the grating).

## Discussion

Optokinetic image stabilization is triggered by slow image drifts but not by fast retinal motion such as occurs during saccades. Here, we identify the inhibitory synaptic circuit primarily responsible for this speed selectivity in mice. It features a familiar amacrine cell type - the VGluT3 cell - in a surprising new role. We show that VGluT3 cells respond well to fast global motion, and that they veto ON DSGC responses to fast motion through feedforward glycinergic inhibition.

Our conclusions are supported by a wealth of convergent evidence. Our pharmacological findings recapitulate earlier evidence in rabbit implicating glycinergic inhibition as the key suppressor of fast motion responses in ON DSGCs. Connectomic analysis shows that just three types of amacrine cells account for nearly all conventional (non-ribbon) synaptic contacts onto ON DSGCs, only one of which - the VGluT3 cell - is glycinergic. We show that the VGluT3 plexus is strongly activated by rapid motion, and that this is true even during global retinal slip, despite the strong suppressive surrounds of VGluT3 receptive fields. We confirm that optogenetic activation of VGluT3 cells evokes glycinergic inhibitory currents in ON DSGCs whereas chemogenetic suppression of VGluT3 output reduces the inhibition of ON DSGCs triggered by rapid motion. Though our data points to feedforward glycinergic inhibition from VGluT3 cells as the main determinant of slow speed preference in ON DSGCs, their speed-tuning profile is undoubtedly shaped to some extent also by their excitatory inputs [15] and by their GABAergic inputs from SACs and widefield amacrine cells.

The output of ON DSGCs to the accessory optic system encodes the local vector of global retinal slip, a precondition for proper image stabilization during head rotation. The vector has two components: direction and speed. The directional coding is well known to derive mainly from a specific amacrine-cell type, the SAC, through spatially asymmetric feedforward GABAergic inhibition of the ON DSGC. We argue that the other vector component is supplied by synaptic input from VGluT3 amacrine cells.

These dual-transmitter interneurons excite some RGCs through glutamate release, while inhibiting others through glycine. An earlier pair of studies reported only excitatory glutamatergic transmission from VGluT3 cells to ON DSGCs [16], not glycinergic inhibition [29]. Though we confirmed some VGluT3-mediated glutamatergic excitation of ON DSGCs, it was present only in a minority of cells and was weaker and more sustained than in the earlier findings. The difference may be partly attributable to our 20-fold dimmer photostimulation stimulus. As far as we are aware, this is the first time any neuronal type has been shown to receive both excitatory and inhibitory input from VGluT3 cells. SACs are also dual-transmitter amacrine cells, exciting DSGCs through nicotinic receptors as well as inhibiting them through GABA. Together, SACs and VGluT3 cells account for more than 92% of the total inhibitory synapses onto ON DSGCs, as shown by our reconstructions. The remaining inhibitory inputs come almost entirely from widefield amacrine cells.

Image stabilization is ubiquitous among animals and crucial for effective vision during self-motion. Optokinetic and optomotor responses are present in all major vertebrate classes and always exhibit preferential activation by slow retinal slip. Many of the genes, cell types, and brain pathways that support such image-stabilizing reflexes are likewise highly conserved in mammals and other vertebrates. Both SACs and VGluT3 cells have been described in higher primate retinas [43–45], so the same two cell types may encode the two components of the global retinal slip vector in the human retina as they do in mice. Genetic defects perturbing this circuit disrupt normal image stabilization in human patients [3].

### VGluT3 cells differentiate the functional signals of ON and ON-OFF DSGCs

It has been known for half a century that ON DSGCs prefer slow speeds while ON-OFF DSGCs prefer faster speeds [8]. The mechanistic basis of this difference has remained obscure. In this report, we show that VGluT3 cells respond vigorously to rapid global motion and use glycinergic feedforward inhibition to suppress ON DSGCs’ response to fast motion. ON-OFF DSGCs, on the other hand, lack VGluT3 inhibition which presumably enables their fast-motion responses. However, VGluT3 cells further differentiate the speed preferences of the two DSGC classes by augmenting fast-motion responses in ON-OFF DSGCs though excitatory glutamatergic synapses.

The two classes of DSGCs have distinct functional roles. ON DSGCs are by far the dominant visual input to the accessory optic system and vestibulocerebellum, which convert slow retinal slip signals into slow optokinetic image-stabilizing eye movements. ON-OFF DSGCs direct their output instead to the superior colliculus and lateral geniculate nucleus, and thus on to the visual cortex. The collicular output allows object motion to trigger gaze or attentional shifts, while the cortical pathway exploits motion information for numerous perceptual computations including figure-ground segmentation, depth perception, and encoding of self-motion. A broad range of speeds are relevant to these midbrain and cortical functions, but for image stabilization, the key motion signal is slow global retinal slip. This specific spatiotemporal pattern of visual reafference informs the brain of a shortfall in the vestibular image-stabilizing reflex, the VOR. If this system were responsive to fast motion, it would conflict with the gaze-shifting network, because it would trigger image stabilization during the fast retinal slip produced by saccades. Silencing the image stabilization system at the retinal level, rather than in downstream brain networks, is energetically favorable since it achieves the same outcome with fewer spikes.

The differing requirements of these two systems in the speed domain is accommodated by the differing speed-tuning profiles of the two classes of DSGCs (ON vs ON-OFF). Our findings now allow us to trace this functional difference back one step further to VGluT3 cells, a interneuron with robust fast-motion-responses and opposing transmitter actions on the two classes: glycinergic inhibition of ON DSGCs, and glutamatergic excitation of ON-OFF DSGCs.

### Are VGluT3 cells the sole source of glycinergic inhibition of ON DSGCs?

Strychnine effectively blocks all glycine receptors on ON DSGCs as shown by the observation that all residual motion-induced inhibition is eliminated by blocking GABA receptors (Fig. 1D). Chemogenetic suppression of VGluT3 cells selectively was less effective in eliminating this feedforward glycinergic inhibition since adding strychnine further reduced it. This may indicate that there are other sources of glycinergic input. The connectomic analysis did identify two synapses from H18 amacrine cells [25, 46] to ON DSGCs. If H18 cells are glycinergic, as suggested by their relatively small, highly branched dendritic arbors, their contributions to ON DSGC inhibition would have been blocked by strychnine but not by our chemogenetic manipulation. However, the transmitter composition of H18 cells is unknown, and they synapse onto ON DSGCs only very rarely. We therefore favor an alternate explanation - that VGluT3 cells are virtually the sole glycinergic input, but that our chemogenetic manipulation of them was only partly effective. This could have occurred for several reasons. When the exogenous receptor was expressed virally, some fraction of VGluT3 cells may not have been transduced. Even when the receptor was presumably expressed ubiquitously through a Cre-dependent genetic cross, application of CNO failed to block the inhibition as completely as strychnine did. This might indicate that the inhibitory K^+^ current evoked by activation of the designer receptor was too weak to suppress all light-evoked synaptic release from VGluT3 terminals. A scaling down of VGluT3 output of this sort could account for the finding that the CNO-sensitive inhibition was a fixed fraction of the total strychnine-sensitive inhibition across the full range of speeds, and that the shapes of the inhibitory current traces were preserved following CNO application (Fig. 5A, B, Supplemental Information and Fig. S4). The alternative model, in which the chemogenetic silencing of VGluT3 cells is incomplete, cannot easily account for such a simple scaling-down of the glycinergic inhibition because the postulated alternative source of inhibition would have to closely match VGluT3 cells in speed tuning. These two models are not exhaustive; others mechanisms might account for the discrepancy between the chemogenetic and strychnine effects. For example, because strychnine blocks glycine receptors globally, it might alter the inhibition measured in ON DSGCs indirectly by its impact on other synaptic circuits.

### New insights into VGluT3 connectivity and function

The synaptic machinery supporting dual glycine/glutamate transmission from VGluT3 cells to their postsynaptic partners remains opaque. The two transmitters could be co-released at the same active zone, with the sign of effect determined mainly by postsynaptic receptor composition. Alternatively, there could be spatially segregated sites of glycine and glutamate release. Our SEM reconstructions provide the first opportunity to compare the ultrastructure of VGluT3 output synapses known to be functionally glycinergic or glutamatergic. Contacts onto known targets of the glutamatergic output (ON-OFF DS; OFF transient alpha; W3/UHD cells) were conventional contacts, not ribbon synapses. Nor were they obviously different in size or form from synapses onto targets known or shown here to be predominantly inhibitory (Suppressed-by-Contrast cells of various forms [29,30,47], ipRGCs [31] and ON DSGCs (present study). Of course, a closer examination or better ultrastructural data might reveal such differences.

The connectomic evidence suggests that the direct influences of VGluT3 cells extend far more widely to other retinal neurons than appreciated in earlier functional surveys. They encompass diverse RGCs and amacrine cells as well as some cone bipolar cells. Other VGluT3 cells are rarely the target of synaptic output from this amacrine cell type, which stands in marked contrast to the extensive mutual inhibition between SACs.

In the absence of obvious ultrastructural correlates of the two sorts of VGluT3 output synapses, we cannot infer whether the previously unknown output targets receive glutamatergic or glycinergic influence from VGluT3 cells. However, our study does offer one small clue. In the presence of glycinergic blockade, optogenetic activation of VGlut3 cells evoked a presumptive GABA-mediated inhibitory current in some ON DSGCs. This suggests that that VGluT3 cells supply glutamatergic excitation to one or both of the GABAergic amacrine cell types that synapse onto ON DSGCs. One of these types, the SACs exhibits no synaptic currents in response to optogenetic activation of VGluT3 cells [16], nor did our SEM analysis turn up more than a very few synaptic contacts from VGluT3 cells to SACs. This leaves the widefield amacrine cells as the likely conduit for this influence. Whatever the circuit responsible, this indirect influence of VGluT3 cells is net inhibitory onto the ON DSGC, just as the direct glycinergic influence is.

Since SEM analysis implied that virtually all types of RGCs receive some VGluT3 input, the role identified here for VGluT3 cells in sculpting the speed-tuning profiles of DSGCs (both ON and ON-OFF) is likely to extend broadly to the retinal output. For RGC types getting excitatory glutamatergic drive from VGluT3 cells, we would expect boosted sensitivity to relatively fast motion. OFF transient alpha cells are one such type and our data confirm their high-speed sensitivity. When the VGluT3 contribution is inhibitory, we expect it to favor responses to slow speeds in the postsynaptic ganglion cells, as for ON DSGCs. Just such a functional influence seems to be detectable in a bistratified medium-field RGC type variously termed ON-delayed [21], Suppressed by Contrast [30,47,48], R-cell [49], or Type 73 [6]. They are ON-type RGCs, but their spiking response to light steps exhibits a characteristic delay which was traced in part to fast feedforward glycinergic inhibition [21]. Ostensibly the same type was inhibited by optogenetic activation of VGluT3 cells [30], and we confirm the synaptic contacts from VGluT3 cells onto this type by SEM. Taken together, these findings imply that this cell type might, like the ON DSGCs, prefer slow speeds. Our survey of speed tuning of various RGC types appears to confirm this (Fig. 1C), since ON-delayed cells were the only RGC type to prefer speeds as slow as those preferred by ON DSGCs.

### The surprising responsiveness of VGluT3 cells to global motion

Ca^2+^ imaging revealed robust responses to global motion in VGluT3 dendrites. This is unexpected because VGluT3 cells are known to be strongly suppressed by visual stimulation of their receptive-fields surrounds [16,20,42]. Responsiveness to global motion also seems at odds with the reported selectivity of these cells for local object motion [20, 42]. We have shown that the strength of surround suppression is context dependent, as is often the case [50, 51]. When the receptive field is probed with a moving grating of various sizes, rather than with flashed spots, the surround appears weaker and the center appears larger ([16], Fig. 6E, F). For gratings instead of spots, both motion and the lack of a sudden and sustained contrast step led to a weaker surround suppression. During retinal slip of a natural visual image, no sudden appearance or large changes in contrast occur, and thus VGluT3 surround suppression should be minimal.

Though we have highlighted here the responsiveness of VGluT3 cells to global motion, they may nonetheless participate in object-motion sensing, as proposed in relation to their glutamatergic excitation of W3 RGCs [20, 42]. Indeed gratings localized to a patch resulted in much stronger responses in VGluT3 dendrites than full-field ones (Fig 6D). The complete suppression measured previously for global motion could well be due to the use of a different stimulus than ours, and specifically the stimulus duration: these studies tested the responses to a short uniform motion of the grating (a 100 µm translation during 0.25-0.5 s). An initial delay in the response relative to the stimulus onset can be seen in our fluorescence traces as well (Fig. 6B) although it could be attributable to voltage-calcium dynamics. Other stimulus parameters (e.g. grating cycle length, orientation, relatively slow speed) could also have an effect.

### Sensitivity of VGluT3 cells to the axis or direction of movement

Virtually every patch of the VGluT3 plexus we tested (95%) showed selectivity for either the axis (orientation) or the direction of grating motion. This was unexpected, and reveals yet another striking parallel with the SACs, whose dendrites are also direction selective. There are also striking differences between the two types in this regard. Though single SAC dendrites prefer somatofugal motion, SAC dendrites of diverse orientations and directional preferences are intermixed at any location in the dendritic plexus, and pooled local Ca^2+^ responses reveal no obvious directional bias at the population level [24]. The VGluT3 plexus, however, shows strong biases in the pooled local signal. An unresolved question is how the directional or orientation preference differs, if at all, between neighboring VGluT3 cells, or within the arbor of a single cell.

If VGluT3 dendrites are orientation/direction selective, can they still suppress fast-motion responses in all four cardinal directions, and thus in all four ON DSGC subtypes? We suggest they do: VGluT3 directional preference exhibited considerable local variability, even when the net preference was strongly skewed. Preferences differed even more across FOVs. Since ON DSGCs have very large dendritic arbors (∼10-12 times larger than the FOV area), they presumably sum inhibition from VGluT3 cells spanning a large region and diverse preferences. Still, the observed bias in VGluT3 motion preferences suggest possible interactions in the optokinetic system between the direction and speed of retinal slip; ON DSGC subtypes preferring some directions may be more strongly suppressed by fast speeds than others.

The connectivity of VGluT3 cells to nearly all RGC types means that their contributions to shaping RGC response properties may extend beyond speed tuning to include selectivity for the direction or axis (orientation) of motion. It will be of interest to assess whether VGluT3 tuning for orientation and direction are linked to their own dendritic asymmetry (as revealed at least in this volume; Fig. 3A-D), similarly to the asymmetric arbors in two RGC types: the ON-OFF DSGCs preferring ventral motion [52] and of JamB-OFF orientation-selective cells [53].

## Methods

### Animals

All procedures were in accordance with the National Institutes of Health guidelines and approved by the Institutional Animal Care and Use Committee at Brown University. Detailed below are the strains of mice used, of adult mice of either sex, 2–8 months old. Wildtype C57BL/6J (Jackson Laboratory); To target ON DSGCs for recording, HoxD10-GFP (GENSAT collection, Tg(Hoxd10-EGFP)LT174Gsat/Mmucd, MMRRC #032065) and Pcdh9-Cre (GENSAT collection, Tg(Pcdh9-Cre)NP276Gsat/Mmucd, MMRRC #036084) were used. The VGluT3-Cre line (The Jackson Laboratory, B6;129S-*Slc17a8^tm1.1(cre)Hze^*/J, #028534) was crossed with Ai32 (Jackson, B6.Cg-*Gt(ROSA)26Sor^tm32(CAG-COP4*H134R/EYFP)Hze^*/J, #024109) for optogenetics, DREADD (Jackson, B6.129-*Gt(ROSA)26Sor^tm1(CAG-CHRM4*,-mCitrine)Ute^*/J, #026219) for chemogenetics, Ai14 (Jackson, B6;129S6-*Gt(ROSA)26Sor^tm14(CAG-tdTomato)Hze^*/J, #007908) for characterization of the VGluT3-Cre mouse, and Ai148 (Jackson, B6.Cg-*Igs7^tm148.1(tetO-GCaMP6f,CAG-tTA2)Hze^*/J, #030328) for Ca^2+^ imaging. For the Müller cell control experiment, GLAST-Cre (Jackson, Tg(Slc1a3-cre/ERT)1Nat/J, #012586) was crossed with Ai32.

### Retinal dissection

Isolation of the retina was performed similarly to [54]. The eyes were removed and immersed in oxygenated Ames medium (95% O2, 5% CO2; Sigma-Aldrich; supplemented with 23 mM NaHCO3 and 10 mM d-glucose). Under dim red light, the globe was cut, and cornea, lens and vitreous humour removed. A relieving ventral cut was made in the eyecup, and the retina was isolated. three more cuts were made in the retina, roughly along the temporal, nasal and dorso-nasal directions, the asymmetry of which was used to disambiguate retinal orientation. The retina was flat-mounted on a polylysine coverslip (Corning, #354086), which was secured in a recording chamber.

### Tamoxifen injections

Tamoxifen (Sigma-Aldrich) was dissolved in corn oil (Sigma-Aldrich) to make 20 mg/ml, sonicated (30 minutes, RT) and placed in hot water (2 hrs, 45 °C) and once homogenous, filtered in a 0.2 µm filter. Tamoxifen was injected IP, 2-2.5 mg per mouse, 3 days in a row. This resulted in dense YFP labeling of Müller glia in GLAST-Cre x Ai32 mice [55]. The animals were used in experiments three weeks after the last tamoxifen injection.

### Electrophysiology

Patch-clamp recordings of isolated flat-mount retina were performed under current-clamp using a Multiclamp 700B amplifier, Digidata 1550 digitizer, and pClamp 10.5 data acquisition software (Molecular Devices; 10 kHz sampling). Pipettes were pulled from thick-walled borosilicate tubing (P-97, Sutter Instruments). Retinas were continuously superfused during experiments with oxygenated Ames’ medium at 32 °C, flow rate ∼5 ml/minute. For cell attached recordings, Ames filled pipettes were used (tip resistance of 4-5 MΩ). For whole cell voltage clamp recordings, pipettes filled with cesium internal solution (In mM: Cs methane sulfonate, 104.7, TEA-Cl, 10, HEPES, 20, EGTA, 10, QX-314, 2, ATP-Mg, 5, GTP-Tris, 0.5, pH 7.3, osmolarity 276 mOsm; All purchased from Sigma-Aldrich) were used (tip resistance of 5.5-6.5MΩ). To isolate excitatory and inhibitory synaptic currents, the recorded cell was held near the reversal potential for inhibition (∼ -65 mV) and excitation (∼ +20 mV), respectively. Application of synaptic blockers to the bath as well as Clozapine-N-Oxide (CNO) was done by switching the perfused medium into medium containing the blocker and waiting for ∼7 minutes. The blockers used were: strychnine (1 µM, Sigma), SR95531 (10 µM, Sigma), L-AP4 (20 µM, Tocris), ACET (10 µM, Tocris), Hexamethonium (100 µM, Sigma) CNQX (20 µM, Tocris), D-AP5 (50 µM, Tocris). In the DREADD experiments, CNO was used to activate the DREADD (1 nM, Sigma).

### Light stimulation

Light stimuli were generated as in [54]. Patterned visual stimuli, synthesized by custom software using Psychophysics Toolbox under Matlab (The MathWorks), were projected (AX325AA, HP) and focused onto the photoreceptor outer segments through the microscope’s condenser. The projected display covered ∼1.5 × 1.5 mm (5.8 µm/pixel). The video projector was modified to use a single UV LED lamp (NC4U134A, Nichia). The LED’s peak wavelength (385 nm) shifted to 395 nm after transmission through a 440 nm short-pass dichroic filter (FF01-440/SP, Semrock), a dichroic mirror (T425lpxr, Chroma), and various reflective neutral density filters (Edmund Optics). The photoisomerization rates used were 10^2^-10^3^ R*/rod/s, and for the used spectrum were similar among rods, M-cones and S-cones. In the beginning of a stimulus sequence, a uniform screen with the stimulus’ mean intensity (gray) was projected for 20-30s for light adaptation.

To identify RGC types, the spike responses of the cell to a 460 µm spot of +0.95 contrast were recorded. To assess the directional tuning of ON DSGCs we used a full-field sinusoidal gratings (cycle = 377 µm, contrast = 0.95, stimulus duration = 5 s, inter-stimulus duration = 3 s at uniform mean grating intensity) drifting in 8 directions in a randomized sequence (drift speed = 226µm/s, 4 repetitions). For speed response curves, the same grating was drifted in the preferred direction (DSGCs), at 7-8 different speeds at a randomized sequence, with 3 repetitions for each speed. For VGluT3 dendrites Ca^2+^ imaging, the same stimulus was used, with the direction of motion chosen arbitrarily. The same was also presented in a circular patch on a gray background in Fig. 6D. In Fig 6E, white spots of different sizes were presented from dark, and the grating stimulus was presented from gray in circular masks of different sizes. In Fig. 6F, gratings and gray spots of different sizes (gray = the mean intensity of the gratings) were both presented from dark, to equalize the global contrast of the two stimuli. To assess the directional tuning of ROIs we used full-field gratings drifting in 8 directions as above, with speeds 1500 or 1900 µm/s.

### Electrophysiology data analysis

Numbers with errors quoted in the text are mean ± standard error of the mean, unless otherwise specified. All data analysis was done using custom written Matlab procedures. Individual peri-stimulus time histograms (PSTH) presented for spikes, or current traces for voltage clamp recordings, were averaged over three trials. Gray traces in Fig. 1B are individual trials. In population response vs. speed data, curves from different cells were normalized by their maximum and averaged. In population data where synaptic blockers or CNO were used (DREADDs), the response curves for each cell were normalized by the maximum of the control curve for that cell, and then curves were averaged over cells. The currents presented for optogenetics (Fig. 4) were recorded at 20kHz, and in Fig. 4B-E and 4J, current traces were averaged in 10ms windows. Current traces in Fig. 4 were averaged over 3-5 trials. Direction and orientation selectivity indices (DSI, OSI) and preferred directions and orientations were calculated from vector sums of responses in different directions of motion [54, 56].

### Immunohistochemistry

Retinas were fixed and counterstained with the following antibodies: Goat anti-ChAT (Choline acetyltransferase; 1:200, Millipore Sigma #AB144); Rabbit anti VGluT3 (1:250, Invitrogen #PA5-85784). Chicken anti GFP (1:1000, Abcam #ab13970) was used to enhance the fluorescence of the Cre-dependent GFP virus. Rabbit anti-HA tag (1: 200, Cell Signaling Technology #3724) was used to stain the HA-tagged hM4Di receptor in the VGluT3 x DREADD mouse.

### Imaging for cell targeting and dendritic morphology

To target fluorescent cells for patch recording, two photon imaging was used (Olympus FV1200MPE BASIC (BX-61WI) microscope, 25×, 1.05 NA water-immersion objective (XLPL25XWMP, Olympus), and an ultrafast pulsed laser (Mai Tai DeepSee HP, Spectra-Physics) tuned to 910 nm. To acquire an image stack, RGCs were filled during electrophysiological recordings with Alexa hydrazide 488 or 594 (100 µM, Invitrogen), and were imaged following the recording, either using the two-photon or the single-photon (confocal) configurations of the two-photon microscope. Tissue in which fluorescent proteins were expressed was often fixed and immunostained (see above), and subsequently imaged on a confocal microscope (Olympus FV3000, UPlan Super Apochromat objectives, 30xS, 1.05 NA, or 60x2S, 1.3 NA) in the Leduc Imaging Facility, Brown University. Confocal and two-photon stacks were processed in Fiji (https://imagej.net/software/fiji), and collapsed using either maximum intensity or maximum standard deviation projections.

### Functional imaging

Imaging of calcium indicator signals were acquired using the aforementioned two-photon microscope and conditions, as has been done previously [54]. The frame rate was 15 Hz. For imaging responses in dendrites, 128 x 256 pixel fields of view were used with a zoom of 4.5-5x (60 µm x120 µm FOVs). Light stimulus presentation was synchronized to the fly-back times in the scanning of the microscope so that they did not interfere with the measured signal.

### Functional imaging data analysis

Functional imaging analysis was done using Fiji and custom written Matlab routines. A standard deviation projection of movies were made, over which ROIs were manually marked over brighter dendrites (Fig 6A), and traces of their area-averaged brightness over time were acquired. For the baseline fluorescence F_0_, we averaged the brightness over 0.5 s before every stimulus presentation. ROIs were chosen for analysis if their time averaged responses surpassed a threshold ΔF/F_0_ (0.3 – 0.6) during at least 6 stimulus presentations out of 24. Fig. 6B-F show curves averaged over responsive ROIs.

### Intraocular injections

Mice were anaesthetized with isoflurane (3% in oxygen; Matrx VIP 3000, Midmark). A viral vector inducing Cre dependent expression of a payload (see below) was injected into the vitreous humour of the right eye through a glass pipette using a microinjector (Picospritzer III, Science Products GmbH). Analgesia (Proparacaine, eye drops) was applied to the eye ∼2 min before the injection, and immediately following the injection (Buprenorphine SR, 0.02 ml, intraperitoneal) to minimize postoperative pain. Mice were then taken off anesthesia, recovered within several minutes, and monitored for 48 hrs following the procedure. Animals were killed and retinas removed 14–21 d later.

### Viruses

pAAV2/2-hSyn-DIO-hM4D(Gi)-mCherry (Addgene #44362, Roth Lab) was injected intraocularly in VGluT3-Cre mice, causing Cre-dependent expression of hM4D(Gi), a modified human muscarinic M4 receptor, that is an inhibitory Designer Receptor Exclusively Activated by Designer Drugs (DREADD). AAV2/2-EF1a-DIO-hChR2(H134R)-EYFP (from UNC vector core, Deisseroth Lab) was injected VGluT3-Cre mice to express ChR2 in a Cre dependent manner, but mostly caused expression in too few of the VGluT3 ACs to drive optogenetic responses in postsynaptic RGCs (Fig. 2E, Supp. Fig. 2D). rAAV2/2-CAG-flex-GFP (UNC vector core) was injected in Pcdh9-Cre mice to target ON DSGCs for recording.

### Optogenetics

Light stimulation to activate ChR2 was generated using a LED light source (Mightex MLS-5500-MK1; LED driver: open-ephys.org, Cyclops) and introduced through the microscope objective and GFP excitation filter cube, resulting in a spectrum peak at 480±10 nm, and illumination over an area of 1 mm in diameter. The light intensity at the sample was 0.9-1.4 nW/µm^2^, the lowest intensity that yielded robust postsynaptic responses in ON DSGCs. The stimulation time was 0.1 s or 1 s for 5 repeats, 4 s between repeats. The intensity and time were optimized for a robust response, while minimizing the rundown of the response over time, and the driving of large bursts of current that were sometimes observed, that were inconsistent over trials or not stimulus-locked.

### Electron microscopy neuronal reconstructions

An existing dataset of retinal sections from a Serial Blockface Electron Microscope (SEM), ‘k0725’ [24] was used. The imaged volume dimensions were 50 x 210 x 260 µm^3^ with the short dimension spanning the IPL and parts of the GCL and INL layers of the retina. The pixel size was 13.2 nm^2^ and the section thickness 26 nm. The images contained intracellular details, e.g. synaptic vesicles. The webKnossos platform (https://webknossos.org) was used for tracing neuronal skeletons and annotating synapses . For the plot of stratification depth (Fig. 3J), the depth of nodes in the skeletons were normalized to the depth of the ON SAC plexus to correct for the tilt and curvature of the tissue in the block [57].

## Acknowledgements

The authors are grateful to Roman Drabchuk for help in intraocular injections and SEM cell tracing, to Rachel Gunderson for careful annotation of the Ca^2+^ imaging movies, and to Dianne Boghossian for technical support. Supported by NIH R01 EY12973.

