## Supplemental Information for "A retinal circuit that vetoes optokinetic responses to fast visual motion"

***Targeting of ON DSGCs for single-cell recording***

We used two-photon illumination to target GFP expressing ON DSGCs in the mouse line HoxD10-GFP, in which all ON DSGCs subtypes and a single subtype of ON-OFF DSGCs express GFP [1]; or the line PCdh9-Cre, in which Cre is expressed by ventral-motion-preferring ON DSGCs [2]. The latter was combined with a Cre-dependent GFP reporter carried by an adeno-associated virus. Alternatively, ON DSGCs were targeted by searching among ganglion cells for their characteristic spiking response to a spot of light projected onto their receptive field center, more sustained and sluggish than for most other RGC types, with a modest onset firing rate [3] (Supp. Fig. S1A). The identity of the cells was further confirmed by their direction selectivity in response to a full-field grating drifting in different directions (Supp. Fig. S1B), and by post-recording imaging of their characteristic dendritic morphology (Supp. Fig. S1C). ON DSGCs are bistratified, with typically one of the largest dendritic fields in the ON layer among RGCs (dendritic diameter 315 ± 11 μm (n=11), in agreement with [1]), and only a handful of short branches in the OFF layer (Supp. Fig S1C).

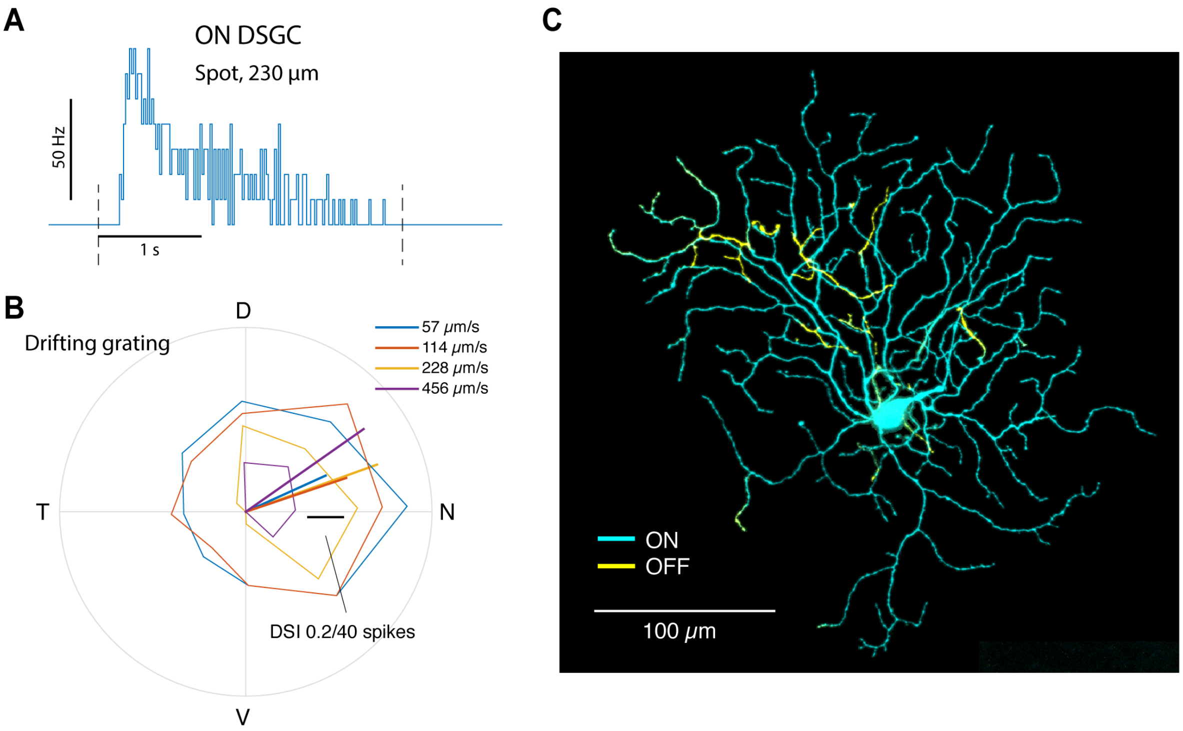


**Supplemental Figure S1. Characteristics of ON DSGCs.** (A) Characteristic response of an ON DSGC to a small bright spot in the center of its receptive field. (B) Directional tuning of ON DSGC responses to full-field drifting gratings, for different grating speed. (C) Dendritic morphology of an ON DSGC. Cyan, dendrites in the ON layer of the inner plexiform layer. Yellow, dendrites in the OFF layer.

##### *Effect of strychnine on inhibition of ON DSGCs*

Strychnine blocks glycine receptors globally in the retina. Using strychnine in the grating experiments (Fig 1D,E in main text), inhibition was reduced by 57%. Following strychnine, the current peaks were dramatically out of phase with respect to the peaks in the control trace, which led us to suspect that strychnine may have introduced unnatural GABA currents. For the slowest speeds this could be seen directly (Fig. 1E, lower left traces). This can occur as a result of blocking of glycine receptors upstream of the RGC, releasing a GABA current in the RGC that would normally be inhibited. If so, then the contribution of glycine to the inhibition is larger than would appear from the data in Fig. 1D (~57%). In support of this, summing the effects of GABA and glycine blockers separately yields a different result than if they were both added together (~60% and 100%, respectively) .

***A VGluT3-Cre mouse line for specific manipulation of VGluT3 amacrine cells***

In order to manipulate VGluT3 cells in a specific manner, we studied Cre-expression in the mouse line VGluT3-IRES2-cre-D that to our knowledge had not been previously used in retinal studies. To this end we crossed the VGluT3-Cre mouse with a tdTomato reporter mouse (Ai14). In retinas of these mice, nearly every VGluT3-immunopositive neuron expressed tdTomato (98.2 ± 0.6%; 9 images, 2 retinas; Sup. Fig S2A). VGluT3-cell dendrites formed a dense plexus concentrated between the ON and OFF SAC plexuses, as revealed by anti-ChAT immunofluorescence (Supp. Fig. S2E). The only other retinal cells brightly labeled with tdTomato were Müller glia, though we occasionally encountered labeled RGCs (~20/mm$^2$) and wide-field amacrine cells with somas in the GCL or INL and straight, sparsely branches processes in the IPL (~2 processes in a 200x200 μm field of view).

To gain optogenetic and chemogenetic access to VGluT3 cells, we crossed the VGluT3-Cre mouse with lines expressing channelrhodopsin (Ai32), or the hM4Di receptor (DREADD), respectively. To verify expression of the reporters in VGluT3 cells in the crossed mice, we used the YFP expressed along with ChR2 in the VGluT3 xAi32 mouse, and an antibody against HA tag was used in VGluT3 xR26 (Sup. Fig

S2B, C). VGluT3 were labeled independently using anti-VGluT3 antibodies, and SACs were labeled using anti-ChAT.

To obtain information on the morphology of the Cre expressing cells in the VGluT3-Cre line, we injected adeno-associated viruses (AAV) expressing GFP or YFP in a Cre dependent manner into the eyes of VGluT3-Cre mice. This often resulted in labeling that was sparse enough for discerning the dendrites of individual cells (Sup. Fig S2A). The morphology of the labeled cells matched that of VGluT3 ACs ([4–6] and SEM data in this study). In addition, the GFP virus resulted in more non-VGluT3 cells expressing GFP than the crossed mouse line VGluT3 xAi14, possibly due to leakiness of the promoter carried by the AAV.

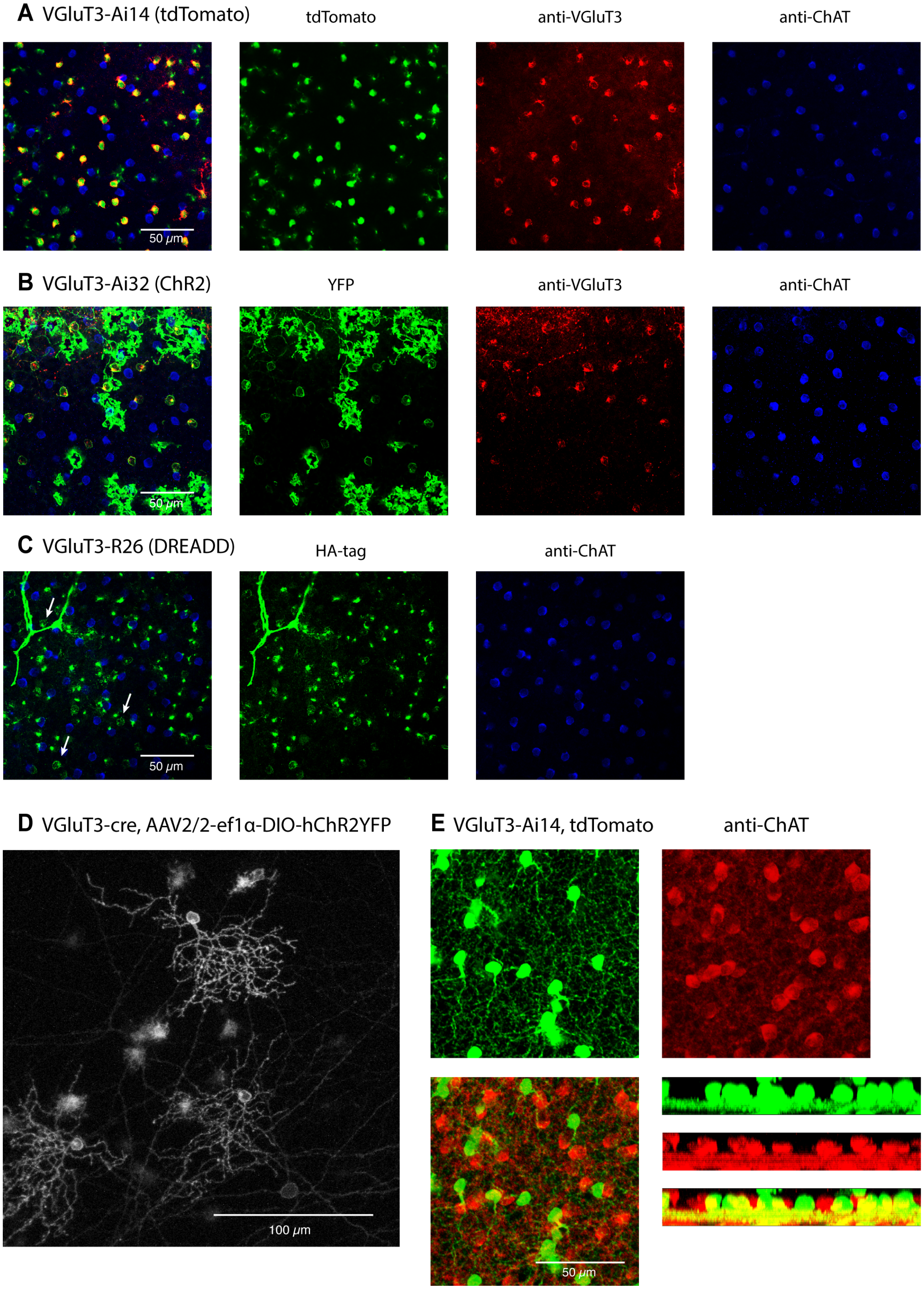
A VGluT3-Ai14 (tdTomato)
tdTomato
anti-VGluT3
anti-ChAT
50 μm
B VGluT3-Ai32 (ChR2)
YFP
anti-VGluT3
anti-ChAT
50 μm
C VGluT3-R26 (DREADD)
HA-tag
anti-ChAT
50 μm
D VGluT3-cre, AAV2/2-ef1α-DIO-hChR2YFP
100 μm
E VGluT3-Ai14, tdTomato
anti-ChAT
50 μm

**Supplemental Figure S2. Characterization of the VGluT3-Cre mouse and its hybrids with reporter mice.** Confocal stacks of VGluT3 and ChAT immunostaining together with (A) tdTomato expressed in VGluT3xAi14. (B) YFP expressed in VGluT3xAi32 (ChR2-YFP) (C) HA tag expressed in VGluT3xR26 (hM4Di-HA). (D) Sparsely labeled VGluT3 cells in a VGluT3-Cre mouse injected with an AAV carrying ChR2-YFP. (E) Top and side views of VGluT3xAi14 at high magnification, showing a VGluT3 dendrite plexus between the two ChAT bands.

##### *Controls for optogenetic studies*

Optogenetically induced currents typically ran down, with smaller responses after repeated stimulation. In a pharmacological experiment, such rundown could masquerade as a drug effect. To assess the extent of the rundown effect, we conducted sham experiments (Fig S3A). Using the standard photoreceptor block but no subsequent drug application, we measured the optogenetically evoked currents at intervals similar to those in an actual pharmacological experiment. Over the time required to complete a real drug trial with two steps of drug application, the peak current dropped by 38 ± 2% (2 cells), but was not eliminated. We conclude that while rundown does contribute to the decreases of the current between steps in these experiments, the pharmacological effects we report are real.

In VGluT3 x Ai32 mice, we observed the expression of the YFP reporter in Müller glia (Fig. S2B). It was therefore important to demonstrate that optogenetic depolarization of Müller glia does not contribute the induced currents in ON DSGCs, especially in light of recent evidence for electrical coupling between Müller glia and amacrine cells [7]. To this end, we crossed a Müller-specific Cre mouse (GLAST-creER) [8] with the Ai32 mouse. In these animals, ON DSGCs exhibited neither excitatory nor inhibitory currents during photostimulation of the Müller cells (Fig. S3B).

The currents induced optogenetically in VGluT3 x Ai32 mice are due to the ChR2 transgene, and not to other light-dependent mechanisms. In control recordings of ON DSGCs in HoxD10 mice without viral or genetic expression of the channelrhodopsin, the optogenetic stimulus evoked no currents (Fig, S3C). This implies that the pharmacological cocktail used in optogenetic experiments to block rod and cone influences on ON DSGCs was largely effective. Rarely, an optogenetic stimulus appeared to trigger large current bursts; these may have resulted from incomplete blocking of rod/cone networks in some cells. Those were mostly inconsistent over trials and had a longer latency compared to persistent optogenetic responses, and were excluded from the data. A persistent ACET-resistant current was recorded in a minority of ON DSGCs, appearing as an OFF response upon termination of the photostimulation (Fig. S3C). Incomplete blockade of light responses tended to be more pronounced in the presence of strychnine.

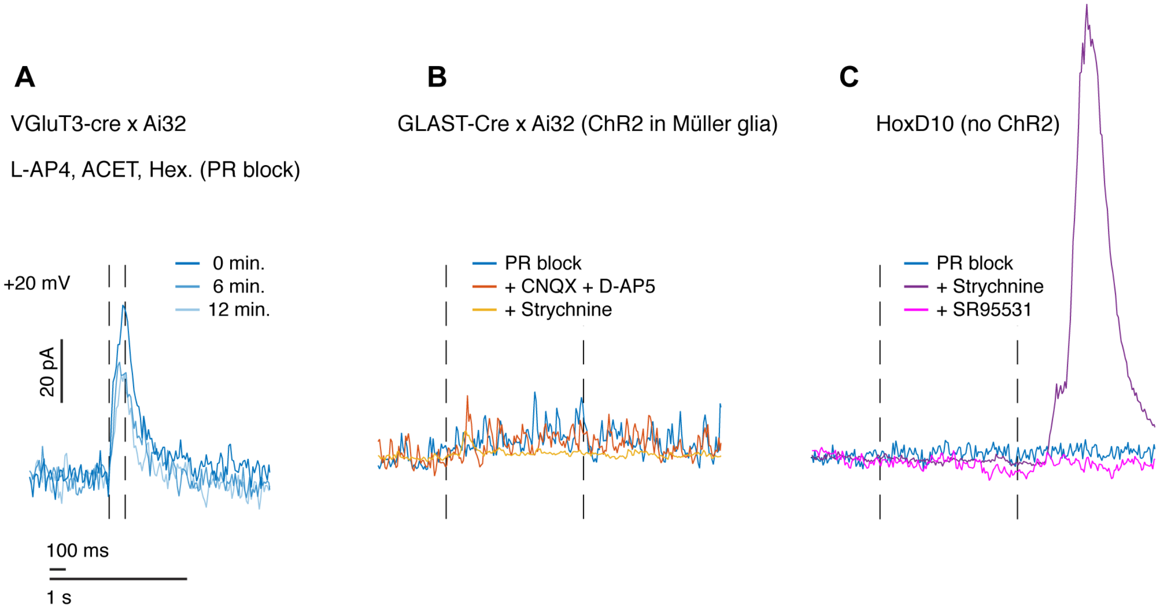


**Supplemental Figure S3. Optogenetics control experiments.** (A) Sham three steps experiment with photo-transduction and acetylcholine block ('PRblock') but without additional blockers added, to test rundown of the optogenetically induced inhibitory current in an ON DSGC. (B) ON DSGC inhibition in a mouse expressing ChR in Müller glia, in response to the LED stimulus, with PR block followed by the addition of glutamate transmission block and strychnine. (C) ON DSGC inhibition in a mouse expressing no ChR, with PR block, followed by the addition of strychnine and SR95531. In A-C, the holding voltage is the excitation reversal potential, and the LED stimulus time is between the dashed lines. Traces are averages over 5 or more trials with a 10 ms window averaging.

***Origin of fast-motion inhibition not blocked by DREADD suppression of VGluT3 cells***

Overall, DREADD-mediated suppression of VGluT3 cells eliminated only about half of the inhibition induced in ON DSGCs by fast motion (Fig 5A, B in main text). The remaining inhibitory current might be supplied by a different type of amacrine cell, but it could also stem from incomplete suppression of VGluT3 cells. Application of the DREADD ligand CNO scaled down of the inhibitory currents while preserving the kinetics (Fig. S4A), and the scaling factor was nearly constant across different grating speeds (Fig S4B). This is most easily explained by incomplete suppression of VGluT3 cells. If a different amacrine type is responsible, it must exhibit temporal kinetics and ON vs OFF sensitivity closely matched to that of VGluT3 cells. The effect of CNO on *ON-OFF DSGC excitation* was not a pure scaling, and was more pronounced at the higher speeds (Fig 5G in main text), in agreement with VGluT3 cells providing only part of the excitation in ON-OFF DSGCs (with the remainder presumably mainly from bipolar cells).

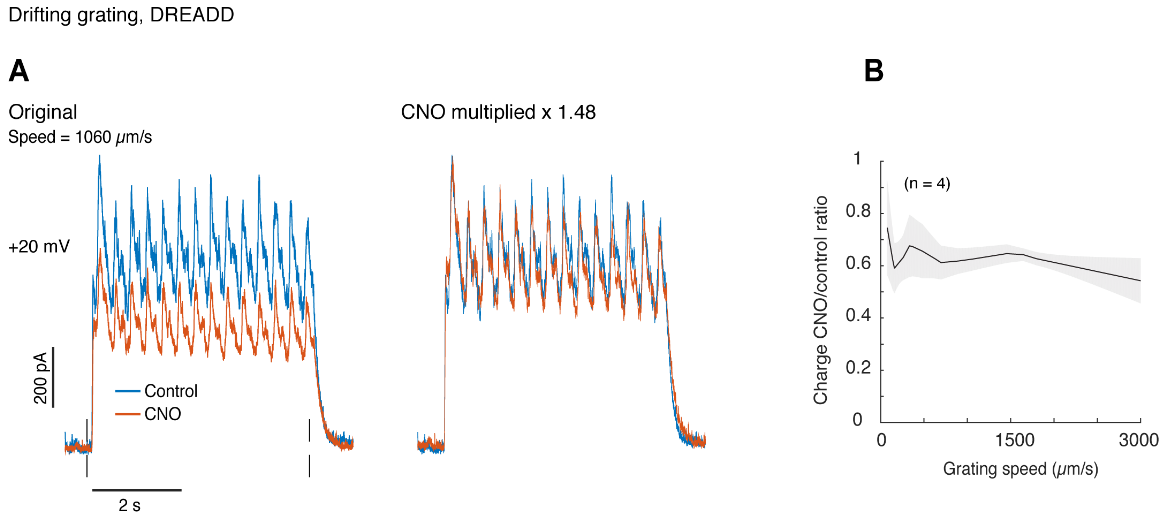


**Supplemental Figure S4. Scaling down of inhibition in ON DSGCs by the DREADD.** (A) Inhibitory current in an ON DSGC before and after the application of CNO (left), and the same currents with the CNO trace multiplied x 1.48, to show high similarity of the curves. The stimulus was a grating drifting in the preferred direction. (B) The ratio of charge transfer before and after CNO, for different grating speeds.

### *SEM data for VGluT3 output synapses*

| | **TYPES** | **EyeWire** [9] | **RGCtypes.org** [3] | **Helmstaedter et al.** [10] | **Other reference** | **Synapses** | **% of cell class (RGC, AC, BC)** |
|---|---|---|---|---|---|---|---|
| **RGCs** | All types | | | | | 537 | 60% of entire sample |
| | M1 | 1ws | M1 | | | 3 | 1 |
| | M2(8) | 9w | M2 | | | 3 | 1 |
| | M2(9) | 9w | M2 | | | 0 | 0 |
| | M4 | 8w | ON alpha | | | 4 | 1 |
| | M5 | 8n/9n? | PixON | 12 | | 0 | 0 |
| | M6 | 91, 915, 81o | M6 | | | 1 | 0 |
| | ON DS | 7id, 7ir, 7iv | ON DS | | | 49 | 9 |
| | ON-OFF DS | 37c,d,r,v | ON-OFF DS | 9 | | 86 | 16 |
| | OIA | 6sw | | | | 20 | 4 |
| | OIB | 6sn | ON transient small RF | | | 15 | 3 |
| | ON transient | 6t | ON transient | | | 11 | 2 |
| | LAMBDA(3a4) | 5so, 51? | HD/LED | 5,10, 11? | | 12 | 2 |
| | LAMBDA(5oi) | 5so, 51? | HD/LED | 5,10, 11? | | 18 | 3 |
| | LAMBDA(5t) | 5so, 51? | HD/LED | 5,10, 11? | | 6 | 1 |
| | W3 | 5ti | UHD | 7 | | 63 | 12 |
| | F-mini ON | 63 | F-mini ON | 6 | | 29 | 5 |
| | OFF trans ALPHA | 4ow | OFF trans alpha | | | 49 | 9 |
| | OMTA | 4on | OFF trans medium RF | | | 34 | 6 |
| | delta | 1wt | OFF sustained alpha | 3 | | 2 | 0 |
| | delta-like | 2aw, 2i, 2o, 3i,3o | OFF sus., OFF medium sus., OFF hOS | 1 | | 3 | 1 |
| | rho | 25 | | 2 | | 4 | 1 |
| | F-mini OFF | 1no, 1ni, 2an, | F-mini OFF | | | 0 | 0 |
| | JAMB | 2w | OFF vOS | | | 1 | 0 |
| | R | 73 | ON delayed | 8 | | 19 | 4 |
| | SBC1 | 81i, 82wi, 82wo | OS | | | 31 | 6 |
| | SBC2 | 72, 82n | OS | | | 11 | 2 |
| | RGC unident. | | | | | 63 | 12 |
| **ACs** | All types | | | | | 337 | 38% of entire sample |
| | A17 | | | 49 | [11] | 0 | 0 |
| | CRH1 | | | 55 | [12, 13, 14] | 1 | 0 |
| | CRH3 | | | 54 | [12] | 0 | 0 |
| | ON SAC | | | 51 | | 2 | 1 |
| | RAC1 | | | | [15] | 0 | 0 |
| | SFE | | | 43 | | 8 | 2 |
| | WF(5i) | | | | | 2 | 1 |
| | WF interChAT | | | | | 25 | 7 |
| | nNOS1 | | | | [12, 16] | 2 | 1 |
| | AII | | | 24 | | 0 | 0 |
| | DAC | | | | | 0 | 0 |
| | H16 | | | 16 | | 0 | 0 |
| | H18 | | | 18 | | 0 | 0 |
| | H19 | | | 17/19 | | 0 | 0 |
| | H21 | | | 21 | | 0 | 0 |
| | H22 | | | 22 | | 0 | 0 |
| | H23 MAC | | | 23 | [7] | 0 | 0 |
| | H36 | | | 36 | | 0 | 0 |
| | H45 | | | 45 | | 5 | 1 |
| | H52 | | | 52 | | 1 | 0 |
| | OFF SAC | | | 33 | | 1 | 0 |
| | SF OFF | | | | | 0 | 0 |
| | TH2 | | | | [17] | 6 | 2 |
| | VGluT3 | | | | | 1 | 0 |
| | VIP-RAC2 | | | | [15] | 7 | 2 |
| | WF ON ChAT + M4 | | | | | 1 | 0 |
| | AC OTHER/ unident. | | | | | 275 | 82 |
| **BCs** | All types | | | | | 22 | 2% of entire sample |
| | GluMI | | | | [18] | 0 | 0 |
| | 1 | | | | | 1 | 5 |
| | 2 | | | | | 0 | 0 |
| | 3a | | | | | 6 | 27 |
| | 3b | | | | | 0 | 0 |
| | 4 | | | | | 4 | 18 |
| | 5o | | | | | 3 | 14 |
| | 5i | | | | | 0 | 0 |
| | 5t | | | | | 3 | 14 |
| | XBC | | | | [3] | 0 | 0 |
| | 6 | | | | | 2 | 9 |
| | 7 | | | | | 0 | 0 |
| | 8 | | | | | 0 | 0 |
| | 9 | | | | | 0 | 0 |
| | RBC | | | | | 2 | 9 |
| | unident. | | | | | 1 | 5 |

**Supplemental Table S5. Output synapses and postsynaptic cells from VGluT3 cells.** n = 896 synapses from 16 VGluT3 cells in the K0725 SEM dataset, organized according to their postsynaptic cells. Since this sample is not random, numbers and percentages may be biased, for example for ON DSGCs. Cell types where matched with nomenclature in three existing datasets, and/or in previous literature.

#### *Supplemental references*


1. Dhande, O.S., Estevez, M.E., Quattrochi, L.E., El-Danaf, R.N., Nguyen, P.L., Berson, D.M., and Huberman, A.D. (2013). Genetic dissection of retinal inputs to brainstem nuclei controlling image stabilization. J. Neurosci. *33*, 17797–17813.
2. Lilley, B.N., Sabbah, S., Hunyara, J.L., Gribble, K.D., Al-Khindi, T., Xiong, J., Wu, Z., Berson, D.M., and Kolodkin, A.L. (2019). Genetic access to neurons in the accessory optic system reveals a role for Sema6A in midbrain circuitry mediating motion perception. J. Comp. Neurol. *527*, 282–296.
3. Goetz, J., Jessen, Z.F., Jacobi, A., Mani, A., Cooler, S., Greer, D., Kadri, S., Segal, J., Shekhar, K., Sanes, J., *et al.* (2021). Unified Classification of Mouse Retinal Ganglion Cells Using Function, Morphology, and Gene Expression. SSRN Electron. J., 1–27.
4. Grimes, W.N., Seal, R.P., Oesch, N., Edwards, R.H., and Diamond, J.S. (2011). Genetic targeting and physiological features of VGLUT3+ amacrine cells. Vis. Neurosci. *28*, 381–392.
5. Haverkamp, S., and Wässle, H. (2004). Characterization of an Amacrine Cell Type of the Mammalian Retina Immunoreactive for Vesicular Glutamate Transporter 3. J. Comp. Neurol. *468*, 251–263.
6. Kim, T., Soto, F., and Kerschensteiner, D. (2015). An excitatory amacrine cell

detects object motion and provides feature-selective input to ganglion cells in the mouse retina. Elife *4*, 1–13.

7. Grimes, W.N., Aytürk, D.G., Hoon, M., Yoshimatsu, T., Gamlin, C., Carrera, D., Ahlquist, R.M., Sabnis, A., Diamond, J.S., Wong, R.O., *et al.* (2020). A high-density narrow-field inhibitory retinal interneuron with direct coupling to Müller glia. J. Neurosci., JN-RM-0199-20.
8. Wang, J., O'Sullivan, M.L., Mukherjee, D., Puñal, V.M., Farsiu, S., and Kay, J.N. (2017). Anatomy and spatial organization of Müller glia in mouse retina. J. Comp. Neurol. *525*, 1759–1777. Available at: https://pubmed.ncbi.nlm.nih.gov/27997986.
9. Bae, J.A., Mu, S., Kim, J.S., Turner, N.L., Tartavull, I., Kemnitz, N., Jordan, C.S., Norton, A.D., Silversmith, W.M., Prentki, R., *et al.* (2018). Digital Museum of Retinal Ganglion Cells with Dense Anatomy and Physiology. Cell *173*, 1293-1306.e19.
10. Helmstaedter, M., Briggman, K.L., Turaga, S.C., Jain, V., Seung, H.S., and Denk, W. (2013). Connectomic reconstruction of the inner plexiform layer in the mouse retina. Nature *500*, 168–174. Available at: https://doi.org/10.1038/nature12346.
11. Grimes, W.N., Zhang, J., Graydon, C.W., Kachar, B., and Diamond, J.S. (2010). Retinal Parallel Processors: More than 100 Independent Microcircuits Operate within a Single Interneuron. Neuron *65*, 873–885. Available at: http://dx.doi.org/10.1016/j.neuron.2010.02.028.
12. Zhu, Y., Xu, J., Hauswirth, W.W., and DeVries, S.H. (2014). Genetically targeted

binary labeling of retinal neurons. J. Neurosci. *34*, 7845–7861.
